# Dynamic coordination of the lever-arm swing of human myosin II in thick filaments on actin

**DOI:** 10.1101/2023.08.30.554051

**Authors:** Hiroki Fukunaga, Takumi Washio, Keisuke Fujita, Masashi Ohmachi, Hiroaki Takagi, Keigo Ikezaki, Toshio Yanagida, Mitsuhiro Iwaki

## Abstract

Muscle myosins work in motor ensembles and must adapt their power stroke in response to mechanical actions by surrounding motors. Understanding the coordination of power strokes is essential for bridging microscopic molecular functions and macroscopic muscle contractions, but the details of this phenomenon remain elusive. Here we used high-speed atomic force microscopy to visualize the individual dynamics (lever-arm swing) of the myosin head bound to actin in DNA origami–based synthetic thick filaments. We observed spatially local lever-arm coordination, and our three-dimensional numerical model explained how mechanical communication between myosins achieved coordination. In a sarcomere model, the local coordination was spatially periodic and propagated toward the contraction direction. We confirmed that a structural mismatch between myosin head spacing (42.8 nm) and the actin helical pitch (37 nm) caused the coordination while improving contraction speed and energy efficiency. Our findings reveal a key physical basis of efficient muscle contraction.

---

Living organisms and their components, from individual molecules to cells and tissues, generate, sense, and react to mechanical forces, engaging in cooperative interactions to perform efficient and adaptive physiological functions^1–6^. Muscle is a representative mechanosensing unit whose functions are attributed to individual myosin motors arranged in a lattice-like structure of thick filaments that interact with actin-based thin filaments in highly structured sarcomeres, the minimal functional units of muscle. A great deal of research efforts has been expended on understanding the molecular mechanisms of muscle contraction^7–14^. Brownian diffusion of the myosin motor domain (head) along actin filament and the lever-arm swing, which involves the rotation of the light chain binding domain (lever-arm) relative to the head coupled with the ATPase cycle^15–18^, is the key dynamic processes causing myosin’s force generation. Recent progress in the study of muscle myosin revealed detailed mechanisms of force generation, including the structure of the actomyosin interface^19, 20^ and two-step structural change of the lever-arm^21, 22^. The force sustained by the actomyosin complex during the cross-bridge cycle likely generates protein deformations that modify the probability of binding, hydrolyzation and unbinding of the ATP molecule with the head. This phenomenon can be seen from a mechanical perspective as a load dependence on the myosin force generation. For example, the detachment rate from actin has been shown to depend exponentially on the external load through Bell’s equation^23–25^.

In myosin ensembles, the lever-arm swing must exhibit adaptive transitions in response to mechanical actions by surrounding motors. Indeed, some previous reports suggested that power strokes occur synchronously among myosin molecules during force generation^26, 27^. Although understanding the cooperative action of the power stroke is important for bridging microscopic molecular function and macroscopic muscle contraction, direct monitoring of individual myosin molecules in muscle has yet to be experimentally achieved, mainly because of the difficulty of accurately labeling biological markers with individual target myosin molecules in intact muscle fibers; in addition, the highly dense structure prevents the analysis of the behavior of individual molecules. Hence, an exciting challenge is to directly and simultaneously visualize the dynamic structural changes of individual myosin molecules in thick filaments and to determine how lever-arm swing dynamics are regulated in motor ensembles.

In this study, we used programmable synthetic thick filaments scaffolded by DNA origami^28–30^, into which recombinant human skeletal myosins (myosin II) was integrated. Unlike conventional synthetic thick filaments composed of purified full-length myosin II^31, 32^, our DNA origami–based thick filaments provide precise control of the motor number and spatial arrangement, which is beneficial for single-molecule observation and quantitative analysis of the motor assembly. We applied high-speed atomic force microscopy (HS-AFM), which can directly visualize dynamic molecular processes^33–35^, to our synthetic thick filaments to simultaneously monitor the behavior of individual myosins. We observed a two-step lever-arm swing, which is consistent with our previous observations^14^, and local coordination in the thick filament. We further developed a coarse-grained three-dimensional model of the interaction between our thick filament and an actin filament to explain how the mechanical correlation between myosins causes locally coordinated power strokes. Additionally, we extended the numerical model to a three-dimensional sarcomere structure in which an actin filament is surrounded by three thick filaments. The physiological geometry induces local and periodic mechanical correlation and coordinated power strokes. We speculate that the structural mismatch between the myosin spacing of 42.8 nm and actin helical pitch of 37 nm^36^ generates the coordinated power strokes. Supporting this notion, in thick filaments with 37-nm myosin spacing, the local coordination disappears. The structural mismatch improved macroscopic mechanical performance, such as contraction speed and energy efficiency, implying that the observed power stroke coordination may be the key to rapid and efficient muscle contraction.

## Results

### Direct observation of human skeletal myosin S1 conformation on actin

We engineered thick filaments using DNA origami and recombinant human skeletal myosin^14^. A 21-base oligonucleotide-labeled myosin IIa subfragment-1 (S1, which contains the head and lever-arm) was attached to linkers (40-nm long two-helix-bundles that mimic myosin II subfragment-2) with the complementary oligonucleotide at 42.8-nm intervals along the thick filament backbone, a setup consistent with the spacing in native thick filaments (Fig. 1a).

**Figure 1.**
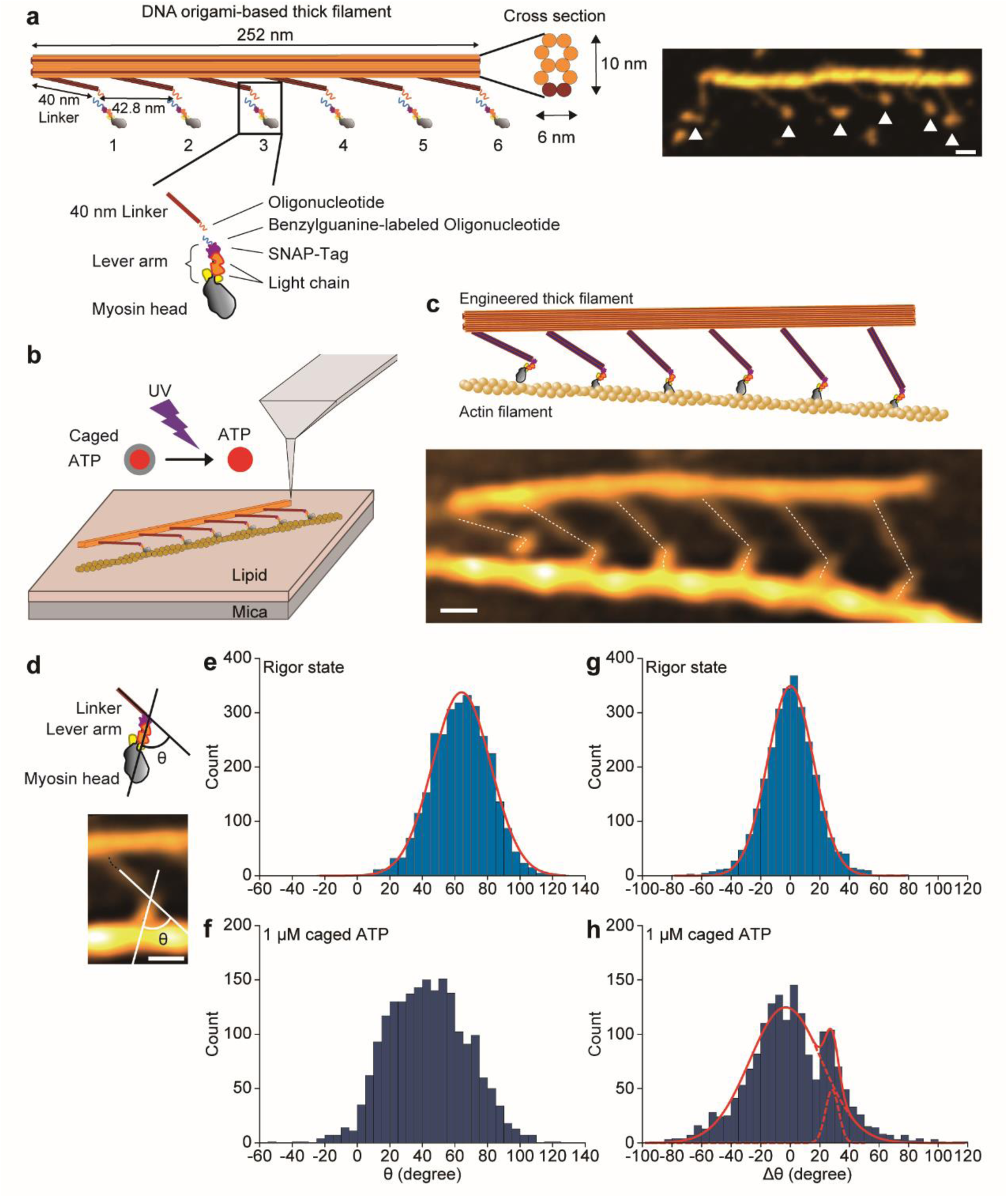
Direct observation of human skeletal myosin S1 conformation on actin. **a** DNA origami–based engineered thick filament. Left, a schematic of the filament. Myosin S1 tagged with SNAP-tag at the C-terminus was attached to the linker through the hybridization of a 21-base oligonucleotide. Right, an AFM image of the DNA origami– based thick filament on mica. Arrowheads indicate myosin heads. Scale bar, 20 nm. **b** Schematic of the AFM observation of the actomyosin complex on lipid bilayers. **c** A complex of the engineered thick filament with actin. Upper, a schematic of the complex; Lower, an AFM image of the actomyosin complex without ATP. White dashed lines indicate the center line of the linker and S1. Scale bar, 20 nm. **d** Definition of the angle (θ) between the linker and the center line of S1. Upper, cartoon showing S1 and the linker; Lower, a typical AFM image. White lines indicate the linear fit of the center line of the linker and S1. Scale bar, 20 nm. **e and f** Angle distribution without ATP (**e**) and with 1 µM caged ATP (**f**). Red solid line indicates fitting of a Gaussian function to the experimental results using a least-squares method with peaks at 64.0 +/- 17.9 nm (mean +/- s.d.). n = 35 and 35 molecules in 7 and 9 actomyosin complexes for e and f, respectively. **g and h** Angle change (Δθ) distribution without ATP (**g**) and with 1 µM caged ATP (**h**). Red solid and broken lines indicate fitting of a single or two Gaussian function to the experimental results using a least-squares method with peaks at 0.19 +/- 15.6 nm (**g**) and -3.1 +/- 24.4 nm and 28.3 ± 4.5 nm (**h**). n = 35 and 35 molecules in 7 and 9 actomyosin complexes for g and h, respectively.

For AFM imaging, DNA-based thick filaments were electrically immobilized on lipid bilayers containing a positively charged lipid on a mica surface (Fig. 1b). To facilitate sideways adsorption of the thick filament onto the bilayer surface, the backbone was designed to have a surface area that was larger on one side than on the other, as shown in the schematic cross-section in Fig. 1a. At low (1 µM caged ATP) or no ATP, we observed single actin filaments bound to myosin heads in the thick filament (Fig. 1b) and the shape of S1 at 2.5 frames/s. We could clearly distinguish the detached and attached states with actin, because the detached myosin fluctuated and became blurry (Supplementary Figure 1 and Supplementary Movie 1), whereas attached myosins showed a globular shape (Fig. 1c). Furthermore, in the attached state without ATP, we confirmed a dominantly kinked structure between the linker and S1, suggesting the post-power stroke conformation (Fig. 1c). We estimated the angle (θ) between the linker and center line of S1 (Fig. 1d; see also Methods) as 64 degrees on average (Fig. 1e). On the other hand, at 1 µM caged ATP, we frequently observed a less kinked structure (∼20 degrees), suggesting the pre-power stroke conformation. Consistently, the distribution of θ in the presence of ATP was broader than that without ATP (Fig. 1f). Regarding the post-power stroke conformation, we frequently detected first and second post-power stroke conformations when a sequential transition occurred (Supplementary Movie 2), consistent with our previous observations^14^. To estimate the conformational change, we confirmed a 28 degree change in θ between adjacent frames in the presence of ATP (Fig. 1g and h). This 28 degrees could not be divided into two peaks (the expected angle change of the pre- to first post-power stroke transition and first post-to second post-power stroke transition); therefore, the angle change for the two transitions are similar, leading us to conclude that θ was roughly ∼20, ∼48 (∼20+28) and ∼76 (∼48+28) degrees at the pre-, first post- and second post-power stroke states, respectively.

### Coordinated power strokes occur locally in the thick filament

Next, to examine the coordinated actions between myosins in our thick filament, we analyzed the conformational state of individual myosins. Figure 2a and b shows an example of the AFM image and possible conformational states for each myosin, respectively. Although we could observe three conformational states on actin (pre-, first post- and second post-power stroke), there were occasions when the states could not be distinguished. Therefore, we measured θ (Fig. 2c, black lines) and its change between adjacent frames (Fig. 2c, bars), then, we defined >20 degree change as power stroke and focused on the power stroke coordination. We frequently observed power strokes in the same frame transitions (Fig. 2b), and their coordination mostly occurred between two or three molecules (∼99%) (Fig. 2d). If power strokes randomly occur between myosins, then coincidental power stroke synchronization should be independent of the distance between them. Therefore, we analyzed the distance dependence of the motors under coordinated motions. Because the number of close myosin pair combinations is larger than the number of distant myosin pair combinations on the filament, we calculated normalized probabilities of the coordinated power strokes as a function of the distance between myosins (separation number) and estimated the distance dependence without bias (Fig. 2e, magenta; see Methods for details). We also computationally generated artificial time series of myosin conformational states, in which the probability of occurrence for each state and transition rates between states agreed with the experimental data (Supplementary Table 1). We regarded each artificial time series as a corresponding myosin and assumed that there was no explicit temporal correlation among state transitions within or between artificial time series. We then calculated the normalized probabilities of the coordinated motions in the artificial time series in the same manner as in the experimental data (Fig. 2e, orange). Because comparisons with random force generation revealed a significant difference, we concluded that coordinated power strokes occurred between adjacent motors in the experimental data.

**Figure 2.**
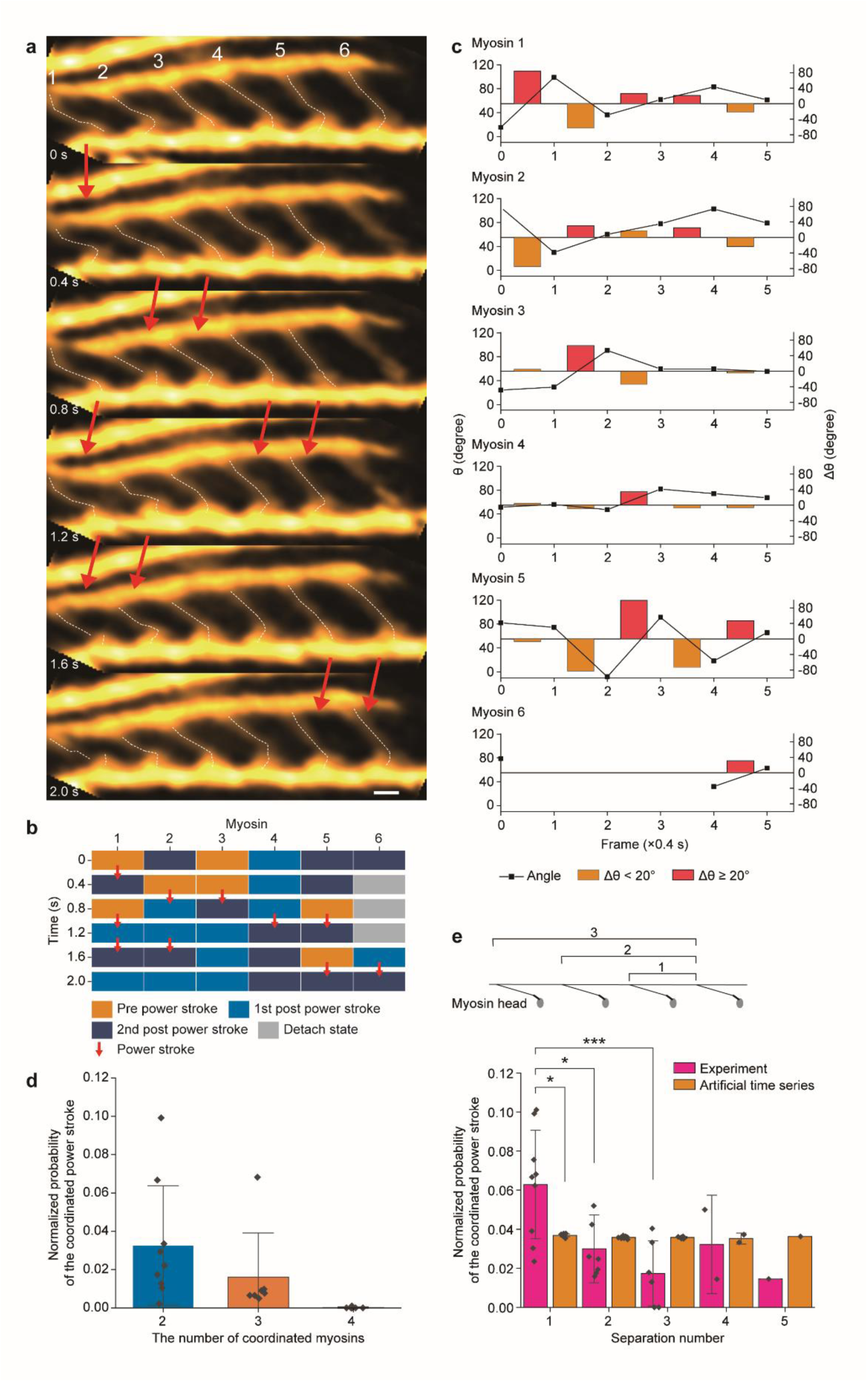
Coordinated power strokes in the engineered thick filaments. **a** Successive AFM images of an actomyosin complex showing the conformational change of S1 in the presence of 1 μM caged ATP. White dashed lines indicate the center line of the linker and S1. Scale bar, 20 nm. **b** Possible conformational states for each myosin in **a**. Red arrows indicate power strokes defined by Δθ > 20 degrees. **c** θ and Δθ between adjacent frames for each myosin in **a**. **d**. Normalized probability of the coordinated power stroke depending on the number of coordinated myosins. n = 68, 23 and 1 coordinated events in 9, 7 and 6 actomyosin complexes for two-motor, three-motor and four-motor coordination, respectively. **e**. Normalized probability of the coordinated power stroke depending on the separation number in the thick filament. In the artificial data, power strokes occurred in a random manner (see Methods). n = 94, 29, 14, 5 and 1 coordinated events in 9, 7, 6, 2 and 1 actomyosin complexes for separation number 1, 2, 3, 4 and 5, respectively, in the experiment. *p < 0.05, ***p < 0.005 (Welch’s two-sided t-test). Error bars indicate standard deviations.

### A coarse-grained model for myosin force generation

To investigate the mechanism of the power stroke coordination in our synthetic actomyosin complex, we developed a three-dimensional coarse-grained elastic network model consisting of our synthetic thick filament interacting with an actin filament. In our previous study, we quantified the flexibility of the linker (Fig. 1a) and tethered myosin heads strongly bound to an actin filament at 36-nm intervals, which reflects the helical structure of the actin filaments (half helical pitch of 36-37 nm^36, 37^), suggesting that steric compatibility between the myosin head and actin regulates the myosin’s binding position on actin^14^. In the coarse-grained model, myosin tethered to a linker (whose flexibility is consistent with experimental data^14^) is represented by nine particles (x_0_–x_8_, Fig. 3a), and the actin filament is represented by double spirals in order to consider the effect of steric compatibility (see also Supplementary Figure 5). The dynamics of the lever-arm swing are represented by the potential function defined on the plane of the two variables: the lever-arm rotation angle 𝜂 and the binding degree of the head with actin 𝜒. These variables are given from the coordinates 𝒙_0_– 𝒙_6_ in the myosin molecule and the coordinates 𝒚_0_– 𝒚_4_ in the actin filament. 𝜒 increases when four virtual particles (𝒙_1_– 𝒙_4_) interact with the points 𝒚_1_– 𝒚_4_ to maintain the appropriate posture at the site of interaction between 𝒙_0_ and 𝒚_0_. Therefore, 𝜒 indicates compatibility between the myosin head and actin helical structure (see Methods for details). The ATPase cycle is reproduced by switching between the two functions 𝜑_𝑃𝑆_ = 𝜑_𝑃𝑆_(𝜂, 𝜒) and 𝜑_𝑅𝑆_ = 𝜑_𝑅𝑆_(𝜂, 𝜒), where 𝜑_𝑃𝑆_ and 𝜑_𝑅𝑆_ simulate the power stroke and the recovery stroke, respectively (Fig. 3b). There are four local minimums on the (𝜂, 𝜒)-plane of 𝜑_𝑃𝑆_, which correspond to the detached state just after the recovery stroke (𝜂 = −90°, χ = 0), the weakly binding state (𝜂 = −90°, χ ≥ 0.45), the pre-power stroke state (𝜂 = −60°, χ ≥ 0.6), the first post-power stroke state (𝜂 = 0°, χ ≥ 0.85), and the second post-power stroke state (𝜂 = 40°, χ ≥ 0.9). An increase of χ for these local minimums with an increase of 𝜂 represents the correlation between the lever-arm swing and the binding strength^20, 23, 38^. These local minimums were chosen so that the experimental HS-AFM results were reproduced well. There are two local minimums on the (𝜂, 𝜒)-plane of 𝜑_𝑅𝑆_ corresponding to the state just after the detachment from the second post-power stroke state (𝜂 = 40°, χ = 1) and the state before the transition to the weakly binding state (𝜂 = −100°, χ = 0). The switching rate is force-dependent, and the dynamics of the lever-arm swing can be modulated by external force. Therefore, mechanical communication between myosins should perturb the dynamics of force generation.

**Figure 3.**
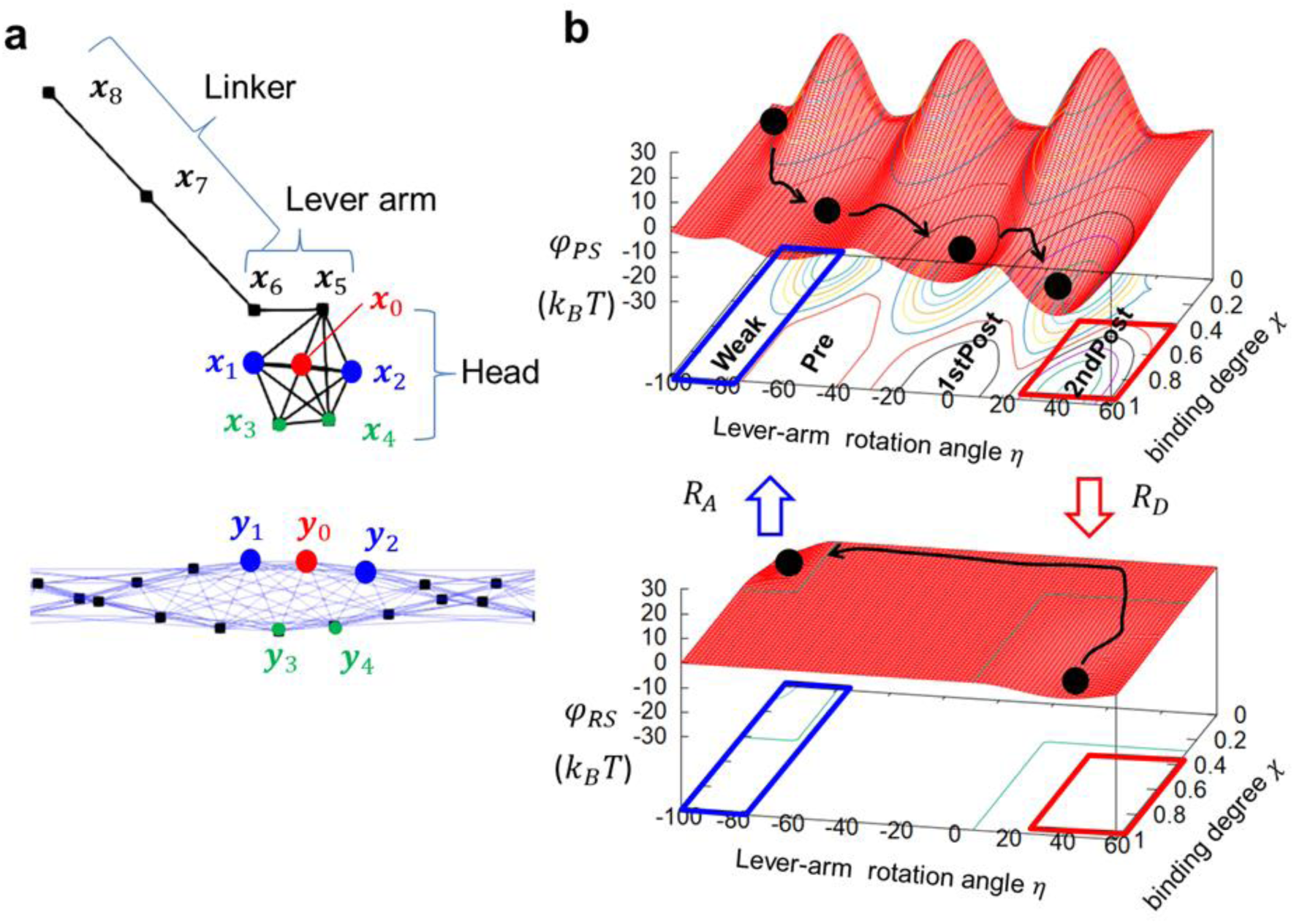
Skeletal structure and potential functions of the numerical model. **a** A myosin molecule is represented by nine particles (𝒙_0_∼𝒙_8_), and the actin filament is represented by double spirals. The lines represent the bonds that hold the basic skeletal structure. In this model, the virtual particles 𝒙_0_∼𝒙_5_, 𝒙_5_∼𝒙_6_ and 𝒙_6_∼𝒙_8_ represent the myosin head, lever arm and linker, respectively. When 𝒙_0_∼𝒙_4_ in the myosin head and 𝒚_0_∼𝒚_4_ in the actin filament are close and compatible with each other, the binding degree 𝜒 increases. **b** The potential functions for the power stroke during the force-producing phase (upper: 𝜑_𝑃𝑆_) and the recovery stroke during the detachment (lower: 𝜑_𝑅𝑆_). These functions are defined on the plane of two variables: the lever-arm rotation angle 𝜂 and the binding degree 𝜒 . As the transition on 𝜑_𝑃𝑆_ progresses, both 𝜂 and 𝜒 increase to represent the lever arm swing and increased affinity with actin, respectively. Then detachment of the myosin head and the recovery stroke smoothly occurs on 𝜑_𝑅𝑆_ (see black ball and arrows). In the ranges surrounded by the red and blue rectangles, switching between the potentials is allowed with the rate constants 𝑅_𝐷_ and 𝑅_𝐴_, respectively.

### Numerical simulation quantitatively reproduces HS-AFM observations

A numerical simulation (Supplementary Movie 3) revealed that the duration of the attached and detached states of myosin with actin (Fig. 4a and b), interval between power strokes (Fig. 4c), actin sliding velocities (Fig. 4d), and distribution of 𝜃 (Fig. 4e) are consistent with those observed in the HS-AFM experiments with 1 µM caged ATP. Furthermore, when we defined power strokes as coordinated if they were observed within 400 ms (time resolution of HS-AFM), we found the power stroke coordination depended on the separation number (Fig. 4f), which is consistent with the experimental result (Fig. 2e). Thus, our three-dimensional coarse-grained model quantitatively reproduced the experimental phenomena.

**Figure 4.**
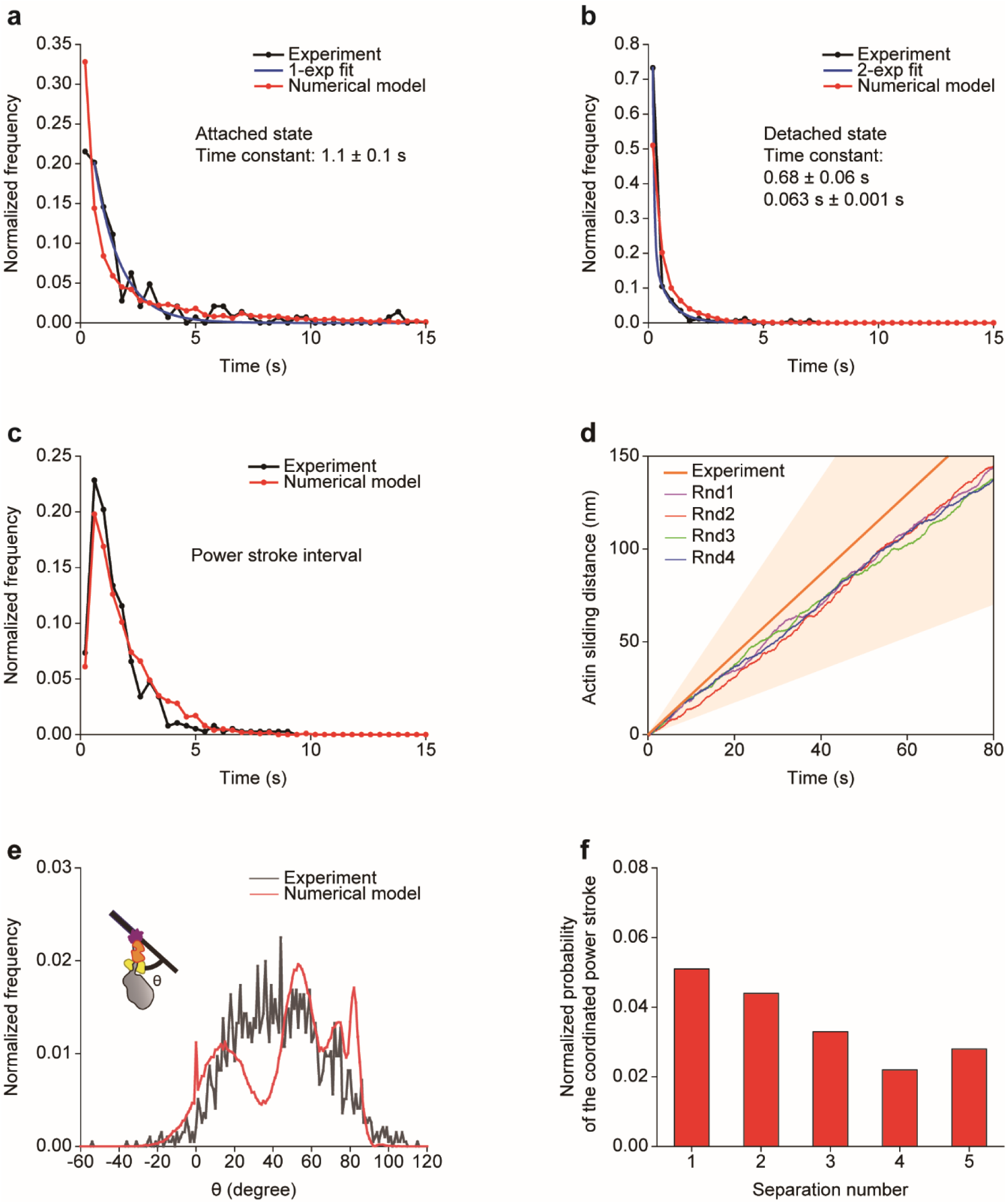
The HS-AFM numerical model reproduces the experimental data. **a–c** Histogram of the duration of the attached state (**a**), detached state (**b**), and interval between power strokes (**c**). Black, red and blue lines indicate experimental results, numerical results and fitting of a single or double-exponential decay curve to the experimental results (1-exp fit or 2-exp fit), respectively. Time constant of the attachment time **(a)** was 1.1 and 1.4 s for the experiment and numerical data, respectively. Time constant of the detachment time **(b)** was estimated to be 0.063 and 0.68 s. The fast time constant was caused by rapid rebinding close to the detached site, and the slow time constant was caused by rebinding to a relatively far site from the detachment site because the actin filament was moving and the detached head could not easily find landing sites in some cases due to the helical structure of the actin. We estimated the mean detachment time as 0.17 s; thus, the duty ratio was 0.87 (1.1 s/(1.1 + 0.17 s)) in the experiment. **d** Displacement of actin sliding for four different sequences of random forces (Rnd1, 2, 3 and 4). The orange line and shadowed area indicate the mean and the standard deviation of the experimental data, respectively. n = 144, 172, 381, and 7 for the experimental data of **a–d**, and n = 319, 584, 562, and 161 for the numerical model of **a–c**, respectively. **e** Distribution of 𝜃 between the linker and lever-arm (see cartoon). Black and red lines indicate experimental and numerical results, respectively. n = 1955 and 83362 for the experimental and numerical data, respectively. **f** Normalized probability of the coordinated power stroke depends on the separation number. The data were taken from simulation results at time intervals [10 s, 80 s] for the four cases with different random force sequences (Rnd1, Rnd2, Rnd3, and Rnd4).

### Mechanical correlation between myosins explains local power stroke coordination

Based on the quantitative model, we investigated how the power stroke of a certain myosin molecule affects other myosins. To this end, we focused on the pulling force of myosin, 𝐹_𝑍_, acting on the actin filament along the spiral. Figure 5a shows the time trajectory of individual myosin conformational states (black lines) in the thick filament and 𝐹_𝑍_ (red lines). 𝐹_𝑍_ fluctuated within 5 pN, a range consistent with the force required for power stroke reversal^39–41^, as observed in HS-AFM experiments^14^, supporting our numerical model. We speculated that the power stroke of a certain myosin mechanically triggers another power stroke; therefore, we focused on a myosin that had completed the second power stroke (“trigger myosin”) and analyzed the force correlation between the trigger myosin and surrounding myosins before the second power stroke (“pre- or first post-power stroke myosin” in Fig. 5b). Figure 5c shows the distance dependency of the force correlation when the downstream myosin is in the second post-power stroke state and the upstream myosin is in the pre- or first post-power stroke state. The trigger myosin generated a larger positive force (i.e., pulling force on actin), whereas the pre- or first post-power stroke myosin generated negative force (i.e., pushing force on actin) for close myosin pairs (Fig. 5d). Because a negative force results in a lower energy requirement for power strokes, the trigger myosin should induce the power stroke for the pre- or first post-power stroke myosin, resulting in coordinated power strokes for close myosin pairs. This tendency was similar when the trigger myosin was upstream (Fig. 5e–g). Therefore, the mechanical correlation effectively explains local power stroke coordination.

**Figure 5.**
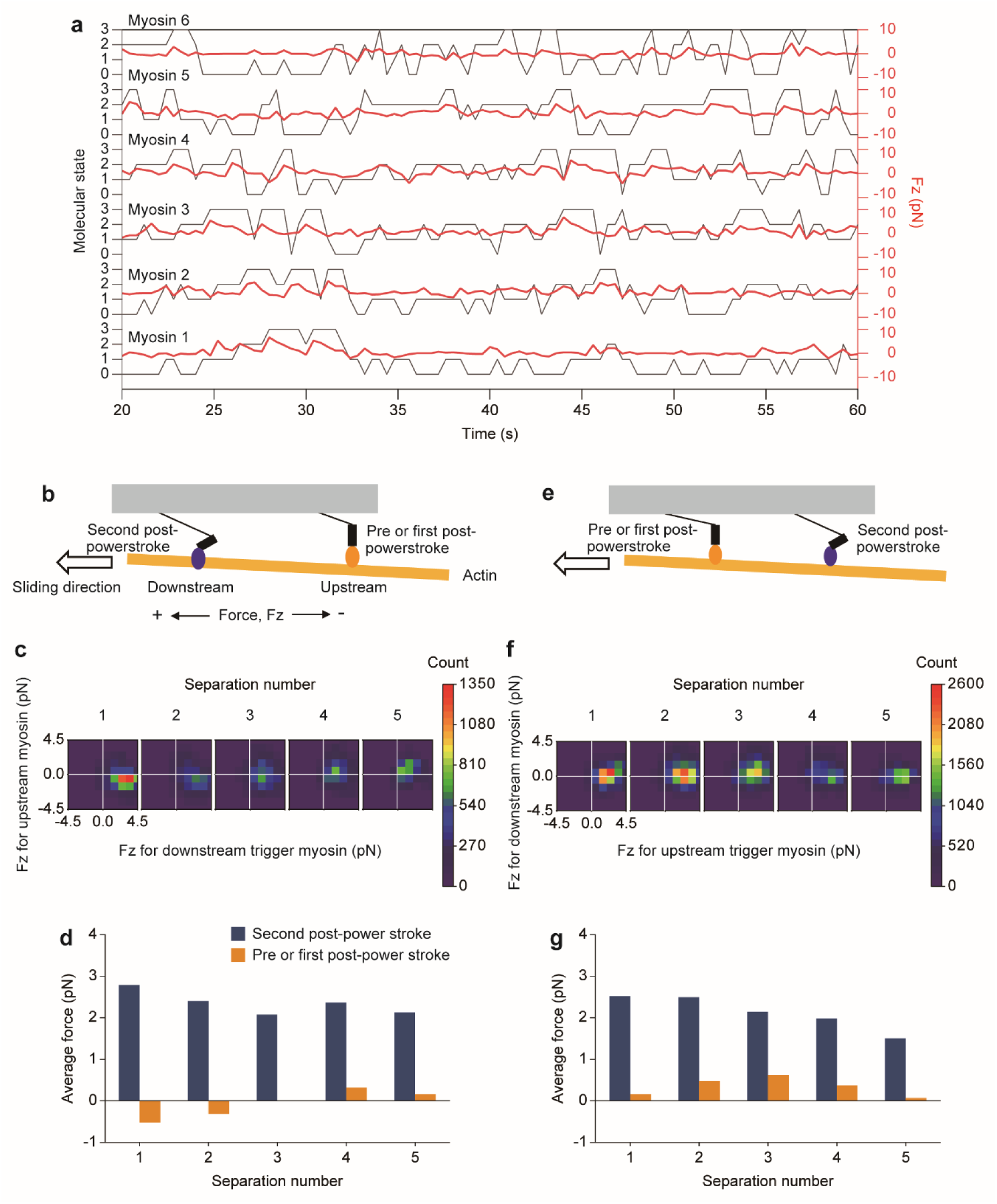
Mechanical correlation between myosins in the HS-AFM numerical model. **a** State transitions (black) and pulling force 𝐹_𝑍_ (red) of the six myosin molecules (Myosin1-6) in the thick filament. The detachment, pre-power stroke state, first post-power stroke state, and second post-power stroke state are represented as 0, 1, 2, and 3 in the left vertical axis, respectively. **b** Schematic of a myosin pair, where a trigger myosin (blue head) and a pre- or first post-power stroke myosin (orange head) are located downstream and upstream, respectively. The actin sliding direction is defined as plus for 𝐹_𝑍_. **c** Contour plots of the force correlation between the downstream trigger myosin (horizontal) and upstream pre- or first post-power stroke myosin (vertical). The plots show individual distances between attached motors. **d** Average force of the trigger myosin (blue) and pre-power stroke myosin (orange) depending on the separation number. **e** Schematic of a myosin pair where a trigger myosin (blue head) and a pre- or first post-power stroke myosin (orange head) are located upstream and downstream, respectively. **f** Contour plots of the force correlation between the upstream trigger myosin (horizontal) and downstream pre- or first post-power stroke myosin (vertical). **g** Average force of trigger myosin (blue) and pre- or first post-power stroke myosin (orange) depending on the separation number.

### Three-dimensional sarcomere structure produces periodic local power stroke coordination

To further investigate the coordinated power strokes in a more physiological geometry, we extended our numerical model to a realistic sarcomere structure in which the actin filament was surrounded by three thick filaments (Fig. 6a and b), each composed of 25 myosin molecules (see also Supplementary Fig. 6b and c, and Supplementary Movie 4). In the simulation, two actin monomers at the right edge of the filament were fixed, because they were connected to the Z-line (Fig. 6c) and were therefore not free to rotate, similar to physiological conditions. Figure 6d shows a kymograph in which individual conformational states of myosin in one thick filament are colored, and time evolution is displayed. Detached and attached (pre-, first post-, and second post-power stroke) states were clustered, and each cluster continuously propagated toward the actin sliding direction at an almost constant speed. The speed was ∼20 nm/s, faster than the sliding velocity of actin (∼3 nm/s at 50 pN load). Here, we used the damping coefficient in the HS-AFM model to describe the actomyosin dynamics (4000 times slower than physiological conditions; see Methods); consequently, the lever arm motion and the transition rate between the active potentials 𝜑_𝑃𝑆_and 𝜑_𝑅𝑆_was also 4000 times slower than physiological conditions (see Supplementary Methods). Therefore, the actual sliding velocity, when the damping coefficient is set to physiological levels, should be ∼12 μm/s (∼3 nm/s × 4000) at a load of 50 pN and ATP concentration of 0.76 mM (190 nM × 4000), in agreement with a previous report^27^. Next, we focused on a myosin pair in the same thick filament (Fig. 6e; the trigger myosin is upstream), and analyzed the average force for the trigger myosin and the pre or first post-power stroke myosin (Fig. 6f). Although both forces were periodic, as the average force of the trigger myosin increased, the force of the pre- or first post-power stroke myosin decreased, implying periodic local power stroke coordination. Supporting this notion, the probability of the coordinated power stroke was local and periodic (Fig. 6g), i.e., “hotspots” for motor coordination. Figure 6h shows the frequency of the observed myosin pair (Fig. 6e), which was periodic, and the high-frequency sites were consistent with that of hotspots. The periodic frequency was probably due to a structural mismatch between myosin head spacing (42.8 nm) and actin helical pitch (37 nm) (Fig. 6b), since the myosin binding site on the actin filaments is helically located, and the head position matches the binding at intervals of ∼6 myosins (i.e. 42.8 nm × 6 = 256.8 nm) (Fig. 6i), consistent with the periodicity of the hotspots (7.2 in Fig. 6h and 6.4 in Fig. 6i). The tendency was similar when the trigger myosin was downstream (Supplementary Fig. 2).

**Figure 6.**
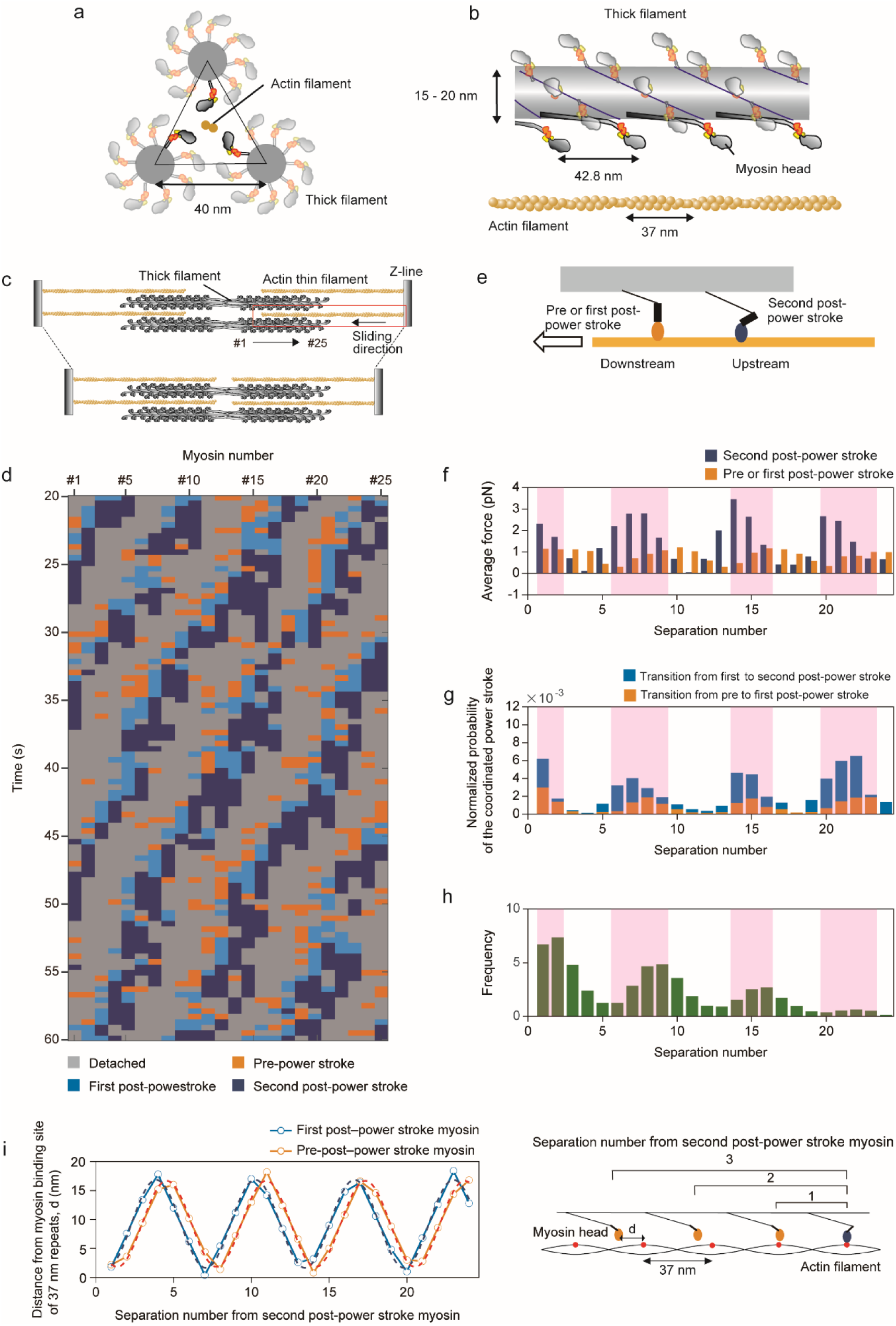
Three-dimensional sarcomere structure produces periodic local power stroke coordination. **a** Schematic of a cross-section of a sarcomere showing three thick filaments with one actin filament in the center. **b** Structures of the thick filament and actin filament. Actin is composed of two protofilaments with a right-handed twist and pseudo-repeat of 37 nm. The myosin heads in the thick filament are arranged in a three-start right-handed helix. The periodicities of the myosin heads and actin filament are different. **c** Schematic of an actin sliding in a sarcomere structure. The actin thin filament is connected with the Z-line and is translocated from right to left. We numbered myosins from left to right (#1 to #25 in the thick filament). The area in the red box was simulated in the sarcomere model. Although one actin filament is surrounded by three thick filaments, only two thick filaments are drawn for visual convenience. **d** A kymograph of the conformational states of 25 myosins in a thick filament shows traveling waves of detached- and attached-state myosin clusters while maintaining cluster size. Data were analyzed at 50 pN load. Load was applied to the actin filament as follows. The right edge was fixed with a stiff spring (10 pN/nm) during initial simulation (0-10 s). Then the spring was disconnected, and the edge was pulled with a constant force in the filament direction. The data analysis was done at steady state at 20-60 s. **e** Schematic of a myosin pair, where a trigger myosin (blue head) and a pre- or first post-power stroke myosin (orange head) are located upstream and downstream, respectively. **f** Average force of a trigger myosin (blue) and pre- or first post-power stroke myosin (orange) depending on the separation number. **g** Normalized probability of the coordinated power stroke depending on the separation number. Orange and light blue bars indicate the fraction for the transition from the pre- to first post-power stroke and first to second post-power stroke, respectively. **h** The frequency of observed myosin pairs depends on the separation number. Pink areas in **f–h** indicate higher cooperative fraction sites (hotspots). All data were analyzed at 50 pN load. **i** Dependency of the distance from myosin binding sites at 37-nm repeats on the separation number from the second post-power stroke myosin. The myosin binding sites at 37-nm repeats are drawn as red circles in the right cartoon. The upstream myosin is in the second post-power stroke state, and the downstream myosin is in the pre- or first post-power stroke state (see right cartoon). The distance *d* was plotted in orange and light blue for downstream pre- and first post-power stroke myosins, respectively. Dashed lines are curves fitted to a sinusoidal function. Both periods are 6.4. We used step sizes of 4 nm for the transitions from the pre- to first post-power stroke and first to second post-power stroke^14^.

### Loss of structural mismatch eliminates hotspot-like coordination

To confirm that the hotspots were caused by the structural mismatch, we analyzed coordinated power strokes for a myosin filament with 37-nm spacing, which matches that of the actin helical pitch (Fig. 7 and Supplementary Movie 5). The kymograph showed that the clusters of detached and attached (pre-, first post-, and second post-power stroke) states were broad, and the propagation became unclear (Fig. 7a). In terms of force correlation, the periodicity disappeared (Fig. 7b), and the probability of the coordinated power stroke exhibited a spatially broad distribution (Fig. 7c), i.e., hotspots for the coordination disappeared. Further, a transition from first post- to second post-power stroke states mainly occurred. Due to the structural matching, the frequency of the observed myosin pair was also spatially homogeneous (Fig. 7d). The tendency was similar if the trigger myosin was downstream (Supplementary Fig. 2). The left-half myosins in the kymograph (far side from Z-line) exhibited a high duty ratio, whereas the right-half myosins experienced long detached periods. This is because rotation of the actin filament is prohibited at the right end of the actin filament, as in the physiological sarcomere structure (the right end of the actin filament is fixed to the Z-line). When the actin filament was translocated, a large rotational motion was observed at the left half of the actin filament (Supplementary Fig. 3), increasing the probability of the rebinding of myosin heads with actin.

**Figure 7.**
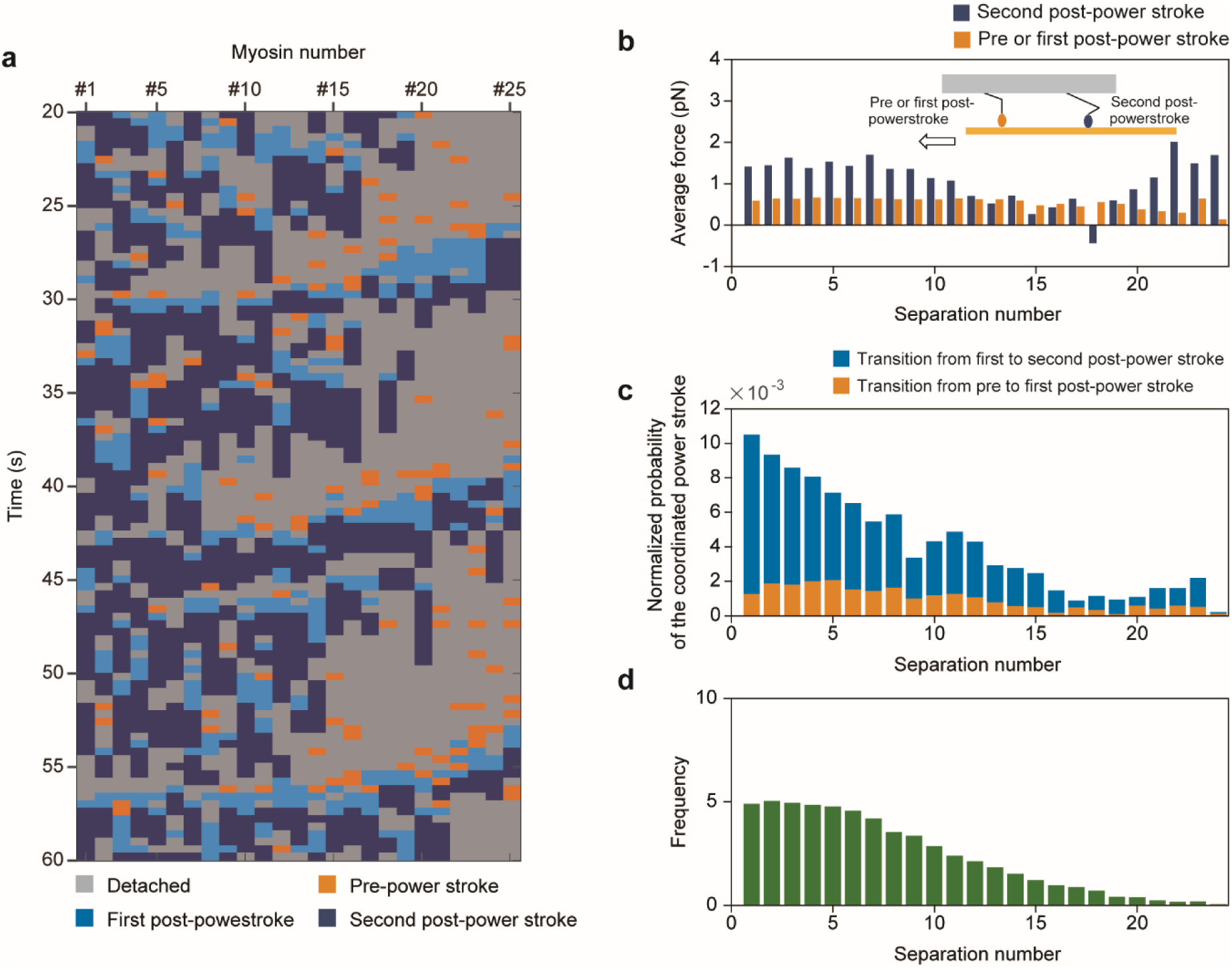
Loss of structural mismatch eliminates hotspot-like coordination. **a** Kymograph of the conformational states of myosins in a thick filament with 37-nm myosin spacing. Data were analyzed at 50 pN load. **b** Average force of trigger myosin (blue) and pre-power stroke myosin (orange) depending on the separation number in a situation in which the trigger myosin is upstream (inset cartoon). **c** Normalized probability of the coordinated power stroke depending on the separation number. Orange and light blue bars indicate the fraction for the transition from pre to first post-power stroke and first to second post-power stroke, respectively. **d** The frequency of observed myosin pairs depends on the separation number. All data were analyzed at 50 pN load.

In addition, we analyzed the mechanical performance of the thick filament with various myosin spacings in the numerical simulation (Fig. 8). The thick filament with 37-nm myosin spacing had the lowest contraction speed without or at low (10 pN) load (Fig. 8a). We then compared the load sensitivity of the contraction speed between spacings of 42.8 nm and 37 nm (Fig. 8b) and found that the contraction speed of the 42.8-nm spacing thick filament was more sensitive to load. Also, the energy conversion efficiency at high load (100 pN) tended to become better at approximately 42.8-nm myosin spacing (Fig. 8c); this spacing showed higher efficiency than 37-nm spacing under every load condition (Fig. 8d).

**Figure 8.**
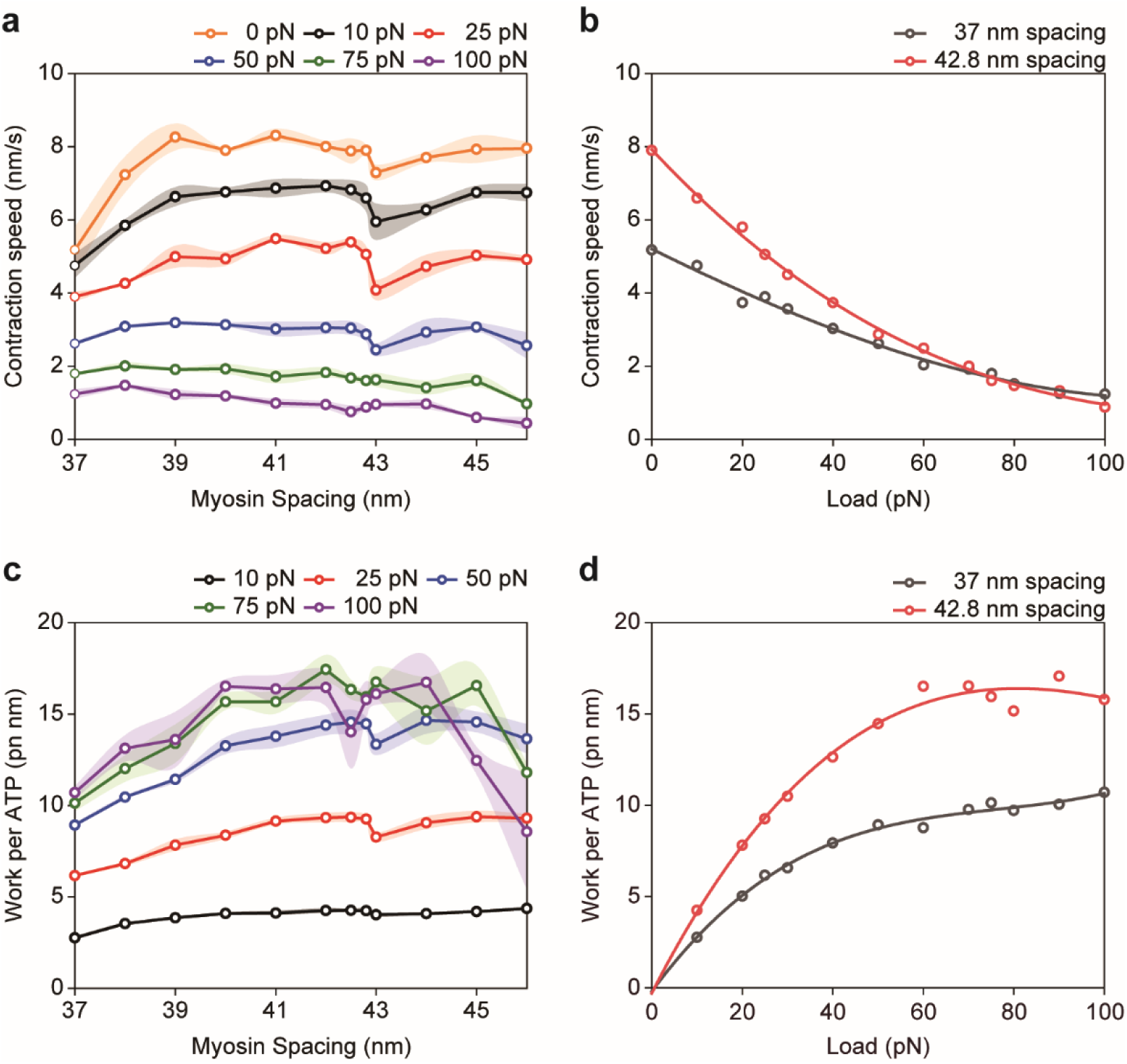
Performance of thick filaments with various myosin spacings. **a** Contraction speed depending on myosin spacing at different loads: 0 (orange), 10 (black), 25 (red), 50 (blue), 75 (green) and 100 pN (purple). The shadowed area indicates standard deviations. N = 3 simulation runs for each condition. **b** Force–velocity relationship for thick filaments with 37-nm spacing (black) and 42.8-nm spacing (red). **c** Work per ATP depending on myosin spacing at 10 (black), 25 (red), 50 (blue), 75 (green) and 100 pN (purple). The shadowed area indicates standard deviations. n = 3 simulation runs for each condition. **D** Relationship between work per ATP and load for thick filaments with 37-nm spacing (black) and 42.8-nm spacing (red). Data plots in **b** and **d** are fitted by a third-order polynomial.

## Discussion

To investigate the power stroke coordination in motor ensembles, we directly visualized the structural dynamics of individual myosin molecules in DNA-based synthetic thick filaments that precisely mimic myosin spacing in the sarcomere. Our results provide insight into the mechanistic details of individual myosins and their coordination in the actomyosin complex. We observed locally coordinated power strokes under our HS-AFM system (Fig. 2e), which could be explained by mechanical communication between myosins. Numerical simulations showed that the coordination strongly depended on the geometry between the thick and actin filaments, i.e., physiological sarcomere geometry with a structural mismatch produced hotspot-like coordinated power strokes (Fig. 6), whereas a structurally matched geometry changed the coordination to a spatially broad distribution (Fig. 7). Furthermore, both detached and attached myosin clusters were continuously propagated toward the actin sliding direction in the physiological geometry. Coupled with the power stroke coordination, the macroscopic performance of the contraction was also altered, and the physiological sarcomere geometry exhibited sensitive load dependency on contraction speed (Fig. 8b) and approximately optimized energy efficiency at high load (Fig. 8c).

How do such propagated coordinated power strokes occur in the physiological sarcomere geometry? In this geometry, highly accessible heads with actin exist at intervals of six myosins (Fig. 6i and Supplementary Fig. 4a). These heads tended to produce coordinated power strokes due to mechanical correlation between myosins (Fig. 6f and g). Because the actin filament is sliding at 3 nm/s at 50 pN load, ∼2 s (that is, ∼6 nm displacement of the actin) is sufficient for the myosin heads in the second post-power stroke state to transition to the detached state (Supplementary Fig. 4b). Other attached myosins transitioned from the pre- to first post-power stroke state or first to second post-power stroke state, and some detached myosins transitioned to the attached state. Thus, attached myosin clusters move toward to the contraction direction with a speed of ∼21.4 nm/s (42.8 nm/∼2 s), consistent with the kymograph (Fig. 6d, Supplementary Fig. 4d). Then, a single head can repeatedly bind to actin with a period of ∼ 12.3 s, because the next actin target comes closer to the head at 37 nm intervals (37 nm/3 nm/s) (Supplementary Fig. 4c). In this calculation, we ignored the rotation of the actin filament during translocation because the rotation speed was slow (∼4.8 degrees in 10 seconds) at the midpoint of the actin filament, which is approximately the same orientation difference for one actin monomer (27.6 degree change in orientation per actin subunit in the actin helical structure) (Supplementary Fig. 3e).

Why does the structural mismatch facilitate efficient contraction speed and energy conversion? When the structural mismatch was dissolved, many myosins can form the strong binding state (Fig. 7a), resulting in a stable actomyosin complex. This geometry produces a large maximum force, and the contraction speed is relatively robust against load (Fig. 8b). However, a coordinated power stroke with a spatially broad distribution will result in a decrease in contraction speed. Furthermore, when we analyzed the distance from the detached myosin head position by ATP binding to the reattached position, the heads in a thick filament with 37-nm myosin spacing rebounded to the positions closest to the detached sites (Fig. 9a), resulting in smaller force generation at pre-power stroke, first post-, and second post-power stroke states (Fig. 9d) and a loss of energy. By contrast, when a structural mismatch exists, accessible myosin binding sites on actin are periodically limited, resulting in a relatively unstable actomyosin complex, reduced maximum force, and higher sensitivity to load. However, most myosin heads can jump and rebind to the 37-nm forward actin binding sites (Fig. 9b), generating greater force (Fig. 9c and d), decreasing energy loss, and obtaining faster contractions.

**Figure 9.**
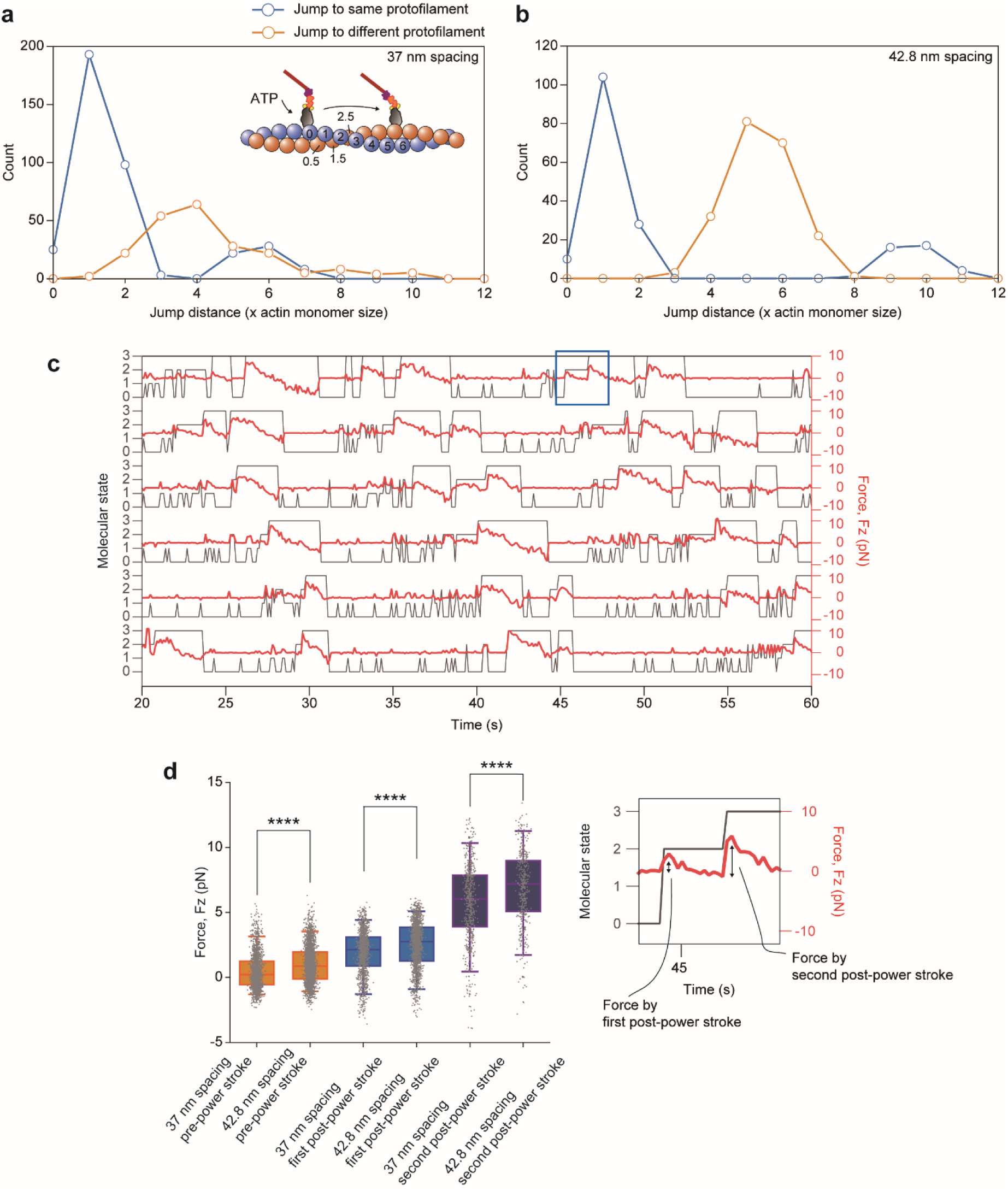
Jump distance and generated force after rebinding. **a** and **b** Jump distances of detached myosin heads for thick filaments with 37-nm spacing **(a)** and 42.8-nm spacing **(b)**. Light blue or orange lines and circles represent the same or different actin protofilament for the rebound position, respectively. Numbering of the jump distance is defined as 1, 2, and 3 for the same protofilament and 0.5, 1.5, and 2.5 for different protofilaments (inset cartoon). **c** State transitions (black) and pulling force 𝐹_𝑍_ (red) of six myosin molecules (myosin #10–15) in the middle of a thick filament with 42.8-nm spacing at 50 pN load. The detached state, pre-power stroke state, first post-power stroke state, and second post-power stroke state are represented as 0, 1, 2, and 3 on the left vertical axis, respectively. **d** Comparison of force between 37 nm and 42.8 nm spacing at the pre-power stroke (orange), first post-power stroke (light blue), and second post-power stroke (blue), respectively. The force just after the state transition was measured (see right graph). n=1720, 2420, 1283, 1516, 618, and 420 for each group (from 75 myosin trajectories), ****p < 0.0001 (Welch’s two-sided t-test). Center line, median; box limits, 1st and 3rd quartiles; whiskers, 5 and 95 percentiles. Right box shows an expanded view of the generated force in the first or second post-power stroke state in the boxed area in **c**. All data were analyzed at 50 pN load.

With regards to the evolution of an animal actuator, by designing a structural mismatch, rapid contraction, higher energy efficiency, and sensitive adaptation to external load may be more important priorities than maximum force and robustness against load. The design principle of muscle we found here will also be useful for artificially constructing nano actuators with desired macroscopic mechanical performance.

## Methods

### DNA origami rod and recombinant proteins with oligonucleotide labeling

DNA origami rods and oligonucleotide-labeled recombinant proteins were prepared as previously described^14^. Briefly, the assembly of the DNA-origami rods was carried out by mixing 50 nM scaffold (p8064, tilibit nanosystems) with 500 nM core staples in folding buffer (5 mM Tris pH 8.0, 1 mM EDTA and 18 mM MgCl_2_) and subjecting the mixture to rapid heating to 80 °C and cooling in single degree steps to 60 °C over 2 h followed by additional cooling in single degree steps to 25 °C over another 72 h. The folded DNA origami rods were purified by glycerol gradient ultracentrifugation^42^. Detailed methods of the myosin construct, expression and purification are described in our previous report^14^. Briefly, the myosin fragment, including the motor domain, essential light chains (ELC) binding domain and regulatory light chains (RLC) binding domain, was cloned from human skeletal muscle myosin IIa cDNA (Kazusa Product ID FXC25901). For oligonucleotide labeling and protein purification, SNAP-tag, FLAG-tag and 6 x His-tag were attached at the C-terminal. These myosin fragments were introduced downstream of the multi cloning site of the pShuttle-CMV vector (Agilent Technologies). Recombinant myosin was expressed using an adenovirus/murine C_2_C_12_ myoblasts system. Recombinant adenoviruses were produced using the AdEasy XL Adenoviral Vector System (Agilent Technologies) and purified using the AdEasy Virus Purification Kit (Agilent Technologies). After the differentiation of C_2_C_12_ cells into myotubes, the cells were infected with the viruses. The myosin was purified using His-tag affinity purification and anion-exchange purification by the AKTA purify system (GE Healthcare). Oligonucleotide labeling reactions were performed just after anion-exchange purification. Amine-modified DNA oligonucleotides (NH_2_/GTGATGTAGGTGGTAGAGGAA) were linked to the benzylguanine (BG; NEB), and BG-oligonuculeotides were labeled with myosin II containing a C-terminal SNAP-tag (NEB) in anion-exchange elution buffer. Oligonucleotide-labeled myosin II was purified by actin filament affinity.

### Preparation of mica-supported lipid bilayers

Lipid vesicles were prepared from a chloroform stock of lipid compounds by mixing them in a glass tube. A typical lipid composition was 1,2-dipalmitoyl-sn-glycero-3-phosphocholine (DPPC; Avanti Polar Lipids 850355) and 1,2-dipalmitoyl-3-trimethylammonium-propane (DPTAP; Avanti Polar Lipids 890870) at a weight ratio of 0.98:0.02. In some cases, the content of the positively charged lipid DPTAP was increased up to 5%. The chloroform was evaporated in a vacuum desiccator for at least 30 min. Lipids were dispersed in Milli-Q water at 1 mg mL^-1^ and stored at -80 °C until use. Supported lipid bilayers (SLBs) were prepared from lipid vesicles via the vesicle-fusion method^43^. Lipid vesicles were diluted to 0.25 mg mL^-1^ in 10 mM MgCl_2_ solution and sonicated to obtain small unilamellar vesicles. SLBs were formed by depositing 2 µL of vesicle solution onto the surface of freshly cleaved mica glued on a glass rod stage (diameter, 1.5 mm) and incubated for 10 min at 60 °C in a humid sealed container to prevent drying of the sample.

### Sample preparation for AFM imaging

DNA rods (10 nM) and myosin-oligonucleotides (1 µM) were incubated for 30 min at 4 °C to form myosin-DNA rod conjugates in advance. After rinsing the lipid bilayer substrate with Milli-Q water and buffer A (10 mM HEPES-NaOH (pH 7.8), 10 mM KCl, 4 mM MgCl_2_, 2 mM EGTA) to remove unadsorbed vesicles, a drop (2 µL) of myosin-DNA rod conjugates in buffer A was deposited on the lipid bilayers for 5 min in a humid hood. After rinsing with buffer A, a drop (2 µL) of actin filaments (1 µM) in buffer A was deposited on the lipid bilayers for 10 min in a humid hood. After rinsing with buffer A containing 1 µM NPE-caged ATP (adenosine 5′-triphosphate, P3-(1-(2-nitrophenyl) ethyl) ester, Invitrogen), the glass rod sample stage was attached to the scanning stage using a wax and immersed in the AFM liquid cell filled with 120 µL buffer A with ATP.

### High-speed AFM imaging

AFM imaging was performed using a HS-AFM system (NanoExplorer, RIBM, Tsukuba, Japan) in tapping mode with a small cantilever (BL-AC10DS-A2, Olympus; resonant frequency, 400∼500 kHz in water; spring constant, 0.1 N/m). Myosin-DNA rod conjugates and actin filaments were weakly adsorbed sideways onto the DPTAP-containing substrate surface. To obtain AFM images of myosin-DNA rod conjugates in the presence of ATP, we typically used a scan area of 300 × 150 nm^2^ with 100 × 50 pixels and imaging rate of 400 ms per frame. UV light irradiation was introduced by a band-path filter (FF01-360/23-25, Semrock) using a mercury lamp light source (U-RFL-T, Olympus) to release caged ATP. All AFM observations were performed at room temperature. The average sliding velocity of actin filaments in the AFM experiments at 1 µM caged ATP was 2.1 +/- 1.3 nm s^-1^ (mean+/-SD). Judging from the actin sliding velocity in our *in vitro* motility assay, the final ATP concentration in the AFM experiments was estimated to be about 190 nM. AFM images were viewed and analyzed using Eagle Software (Ibis_1.0.0, RIBM, Tsukuba, Japan) and ImageJ (National Institutes of Health). We applied to each AFM image a Gaussian filter to remove spike noises. Then, we subtracted background and adjusted contrast for clarity using ImageJ 1.50i.

### Analysis of actin sliding velocity

To estimate the actin sliding velocity, we tracked the end of the actin filament in each frame by Image J, and the distance from the initial point was plotted as a function of time. Then, the regression line was obtained, and we calculated the mean and standard deviation from the slopes.

### Angle between linker and S1

To analyze the angle between the linker and S1, we referred to a previous study^44^. We estimated the center of mass of the x, y coordinates of S1 and linker using multi point tool featured in ImageJ. We used the x, y coordinates to estimate the center line of S1 and linker and then measured the angle between the two lines according to equation:

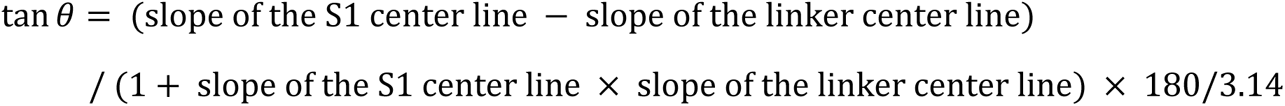

where slope of the S1 and the linker center line = 𝛥𝑦/𝛥𝑥, respectively.

### Probability of coordinated myosin number

The normalized probability of coordinated motion depending on the coordinated myosin number was calculated by dividing the observed number of coordinated motions by the total number of possible combinations depending on the number of coordinated myosins. The total number of possible combinations was calculated from the number of possible coordinated motions in a frame multiplied by the number of video frames. (For example, the possible combinations for two myosin coordination in a frame are _6_C_2_ = 15.)

### Distance dependency of coordinated motions

The normalized probability of coordinated motion was calculated by dividing the observed number of coordinated motions by the total number of possible combinations depending on the coordinated distance. The total number of possible combinations was calculated from the number of possible coordinated motions in a frame multiplied by the number of video frames. (For example, possible combinations of coordinated motions between neighbors in a frame are 1-2, 2-3, 3-4, 4-5 and 5-6 (motor number on the thick filament), so the number of possible combinations is 5. On the other hand, a possible combination of coordinated motions between motors that are five molecules apart in a frame is only 1-6 (motor number), so the number of possible combinations is 1.) Furthermore, we computationally generated artificial time series of myosin conformational states so that the probability of occurrence for each state and transition rates between states agreed with the experimental data. The initial conformational state for each myosin was also stochastically determined based on the probability of occurrence for each state, and then these conformational states individually transitioned frame by frame according to the transition rates. We assumed no explicit temporal correlation among state transitions within or between artificial time series. The normalized probability of the coordinated motion occurring in the artificial time series was calculated in the same manner as the experimental data. We repeated this process 1000 times to estimate the average and SD of the probability, and statistically evaluated the experimental data.

### Numerical Model

To simulate the dynamical motion of myosin molecules and the actin filament in three-dimensional space, we developed a three-dimensional coarse-grained molecular dynamics model governed by the overdamped Langevin equation:

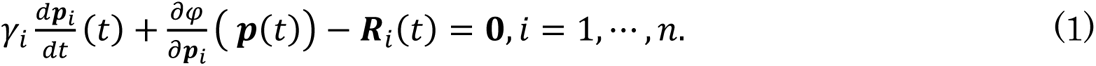

Here, 𝛾_𝑖_ is the damping coefficients, 𝑹_𝑖_ = (𝑅_𝑥,𝑖_, 𝑅_𝑦,𝑖_, 𝑅_𝑧,𝑖_) is a white and Gaussian random force with the mean 〈𝑅_𝑥,𝑖_〉 = 〈𝑅_𝑦,𝑖_〉 = 〈𝑅_𝑧,𝑖_〉 = 0 and the variance

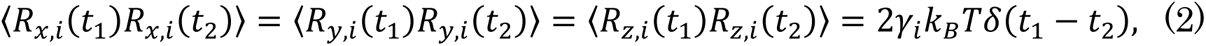

where Boltzmann’s constant is 𝑘_𝐵_, and the temperature is 𝑇. In our simulations, we used 𝑘_𝐵_𝑇=4.128 pN·nm assuming room temperature 𝑇=26 ℃. The potential energy denoted by 𝜑 is given as a sum of the two functions of the particle positions 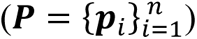:

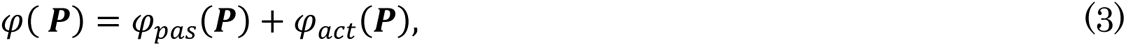

where 𝜑_𝑝𝑎𝑠_ is the passive potential holding the skeletal structure of the myosin molecules and the actin filament, and 𝜑_𝑎𝑐𝑡_ is the active potential which drives the lever-arm rotation and the binding of the myosin head with the actin filament. This active potential is switched between the two functions, 𝜑_𝑃𝑆_ and 𝜑_𝑅𝑆_, under the stochastic rule: 𝜑_𝑃𝑆_ facilitates the forward power stroke during the force producing phase, and 𝜑_𝑅𝑆_ facilitates the recovery stroke that returns the lever-arm position to the pre-power stroke state during the detachment^45^.

As shown in the Results section, the cross-bridge cycling in the HS-AFM environment is extremely slower than that in the physiological condition. In our numerical model, we assumed that this slowdown originates from a higher damping coefficient 𝛾_𝑖_ compared to that in the physiological condition. In the Langevin equation, the damping effect is assumed to originate from the collisions with solvent molecules. In the physiological condition, the solvent molecules almost entirely consist of water molecules. However, in the HS-AFM condition, the dominant source of damping may be collisions with the lipid particles, because the actomyosin molecules are absorbed onto the positively-charged lipid bilayer on mica. Since the lipid particles are much bigger and heavier than water molecules, the damping coefficients can be much larger than in the physiological condition. In our numerical model, we applied a damping coefficient 4000 times larger than that in the physiological condition to fit the dwell times of each conformational state and actin sliding velocity (Fig. 4). Note that the multiplications of the damping coefficient by 𝛼_𝑑𝑎𝑚𝑝_ and switching rate constants between the active potentials 𝜑_𝑃𝑆_ and 𝜑_𝑅𝑆_ by 1/𝛼_𝑑𝑎𝑚𝑝_ result in slowing down the motion by 1⁄𝛼_𝑑𝑎𝑚𝑝_ (see Supplementary Methods A) according to Equation (1) with the random force defined in Equation (2). Furthermore, to adjust the bending behavior of the actin filament, as observed in the experiment (data not shown), the damping coefficients for the particles in the actin filament were multiplied by twenty. This additional damping is not reflected in the magnitude of random force 𝑹_𝑖_ in Equation (2), because we assume that it comes from the attraction between the negatively charged actin monomers and positively charged lipid particles.

In the following, the key factors of the potential model are described. The details and the model parameters are given in Supplemental Methods. In our model, the actin filament is represented by the double spirals of the binding points. Each particle corresponds to an actin monomer which is lined up along one of the two spirals. The half helical pitch of the actin filament was assumed to be 36 nm without load (HS-AFM model) and 37 nm with load (sarcomere model)^36^. The stiffness of the actin filament is given by the harmonic potentials associated with the bond length between neighboring particles, as depicted by the thin lines in Fig. 3a, and the harmonic potential for the dihedral angle between the two neighboring triangles (Fig. 3a). The stiffness parameters were adjusted so that the persistence length of an actin filament (15 μm)^46^ can be reproduced. The myosin molecule is represented by nine particles (Fig. 3a). 𝒙_0_ is the binding point with the actin filament, and the line segment 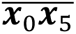 corresponds to the myosin head. The line segment 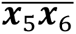 is the lever-arm that rotates around 𝒙_5_ during the power stroke and the recovery stroke. The linker consists of two line segments, 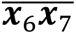 and 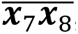 where 𝒙_8_ is connected to the thick filament backbone. A harmonic potential for the bond angle between 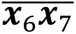 and 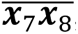 was applied to represent the bending stiffness of the linker^14^. The four points 𝒙_1_, 𝒙_2_, 𝒙_3_ and 𝒙_4_ are virtual particles whose positions relative to the central binding point 𝒙_0_ were determined according to the arrangements of the five binding points 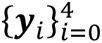 in the actin filament. The point 𝒙_0_ is placed 90° from the top position of the cylinder of the actin filament.

A key distinguishing feature of our model is the active potential energy, which represents the correlation between the lever-arm swing and the binding strength with the actin filament. Thus, it is given as a function of two variables, 𝜂 and 𝜒, where 𝜂 is the lever-arm rotation angle, and 𝜒 represents the binding degree of the myosin head with the actin filament. 𝜂 is defined as

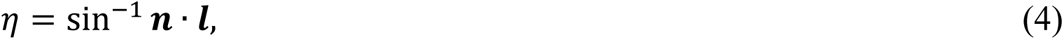

where the unit vector 𝒏 is the direction vector along the spiral of the actin filament given by 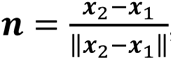, and the unit vector 𝒍 is the direction vector of the lever-arm given by 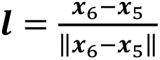. The binding degree 𝜒 is defined as

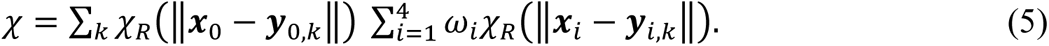

where 𝜒_𝑅_ is given as

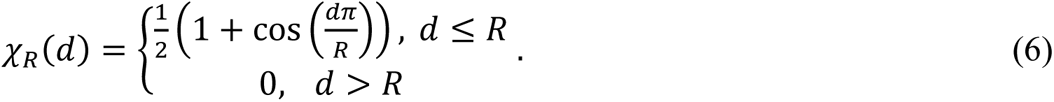

The summation over 𝑘 is taken through all actin monomers 𝒚_0,𝑘_, for which 𝒚_1,𝑘_ and 𝒚_2,𝑘_ are two neighboring points in the same spiral, and 𝒚_3,𝑘_ and 𝒚_4,𝑘_ are two neighboring points in the counter spiral. By representing the binding degree with the product as in Equation (5), only the myosin heads in proper posture relative to the actin filament are allowed to bind. In this study, 𝜔_1_=𝜔_2_=𝜔_3_=𝜔_4_=0.25 were used. By applying larger weights to the virtual points that correspond to the counter spiral, unnatural bindings with the actin filament can be prohibited. As the effective radius, 𝑅=5 nm was chosen. With this choice, even if a non-zero value of the product is given for two indices of 𝑘, one of the products is very close to zero.

Following the above definition of 𝜂 and 𝜒, the ATPase cycle is reproduced by switching between the two functions 𝜑_𝑃𝑆_ = 𝜑_𝑃𝑆_(𝜂, 𝜒) and 𝜑_𝑅𝑆_ = 𝜑_𝑅𝑆_(𝜂, 𝜒), where 𝜑_𝑃𝑆_and 𝜑_𝑅𝑆_ were applied to simulate the power stroke and the recovery-stroke, respectively (Fig. 3b, Supplementary Methods G). The force-dependent switching rate^27^ from 𝜑_𝑃𝑆_ to 𝜑_𝑅𝑆_ is given as

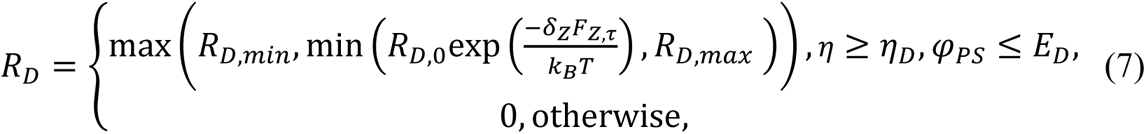

where 𝐹_𝑍,𝜏_ (simply denoted as 𝐹_𝑍_ in the main text) is the filtered value with the time constant τ of the pulling force imposed on the actin filament along the spiral:

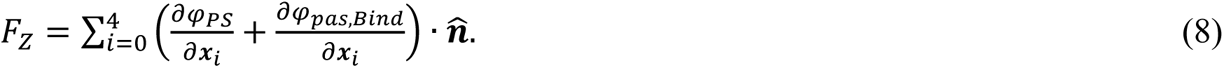

Here, 𝜑_𝑝𝑎𝑠,𝐵𝑖𝑛𝑑_ is the passive binding potential (See Supplemental Methods H), and 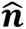 is the unit vector directing the actin filament.

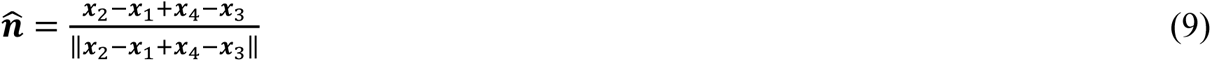

This switching may correspond to either the dissociation of an ADP molecule or the binding of an ATP molecule. We assumed that the dissociation rate decreases as the pulling force increases following Bell’s approximation47. In our model, the difference between 𝜑𝑅𝑆 and 𝜑𝑃𝑆 on the dissociation region is about 25 𝑘_𝐵_𝑇, which is nearly equal to the ATP hydrolysis energy. The switching rate from 𝜑_𝑅𝑆_ to 𝜑_𝑃𝑆_ is given as

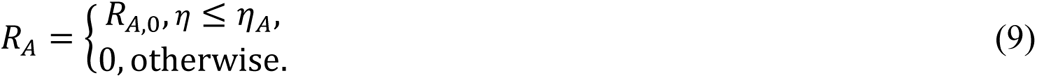

This means that the switching is applied after the recovery stroke is accomplished. The parameters 𝑅_𝐷,0_ =2[1/s], 𝑅_𝐷,𝑚𝑖𝑛_ =0.1[1/s], 𝑅_𝐷,𝑚𝑎𝑥_ =10[1/s], 𝜂_𝐷_ = 30°, 𝐸_𝐷_ =−15.5 𝑘_𝐵_𝑇, 𝛿_𝑍_=2.12nm, and 𝑅_𝐴,0_=100[1/s], 𝜂_𝐴_=−80° were adopted in the HS-AFM model, while the factor 0.25 was multiplied to the rate constants in the sarcomere model to attain better performance.

In our model, the ATPase cycle is completely characterized by the two potential functions 𝜑_𝑃𝑆_ and 𝜑_𝑅𝑆_ and the switching rates 𝑅_𝐴_ and 𝑅_𝐷_. Among them, 𝜑_𝑃𝑆_ is the key factor and represents the correlation between the lever-arm swing and binding strength during the force producing phase. There are four local minimums on the (𝜂, 𝜒)-plane corresponding to the detached state just after the recovery stroke (𝜂 = −90°, χ = 0, 𝜑_𝑃𝑆_ ≈ −4 𝑘_𝐵_𝑇), the weakly binding state (𝜂 = −90°, χ ≥ 0.45, 𝜑_𝑃𝑆_ ≈ −8 𝑘_𝐵_𝑇), the pre-power stroke state (𝜂 = −60°, χ ≥ 0.6, 𝜑_𝑃𝑆_ ≈ −15 𝑘_𝐵_𝑇), the first post-power stroke state (𝜂 = 0°, χ ≥ 0.85, 𝜑_𝑃𝑆_ ≈ −18.5 𝑘_𝐵_𝑇), and the second post-power stroke state (𝜂 = 40°, χ ≥ 0.9, 𝜑_𝑃𝑆_ ≈ −29 𝑘 _𝐵_𝑇). One of the interesting points is that the energy difference between the pre-power stroke state and first post-power stroke state is 3.5𝑘_𝐵_𝑇, while the difference between the first post-power stroke state and second post-power stroke state is 10.5𝑘_𝐵_𝑇. This seems to be inconsistent with the ratio of the lever-arm swing angles of the first and second post power strokes (60°: 40°). The balance between the energies and the lever-arm rotation degrees determines the dependence of the lever-arm swing on the mechanical load. Another interesting point is that the binding energy at the pre-power stroke state (15𝑘_𝐵_𝑇) is more than the half of the binding energy after the second power stroke state (29𝑘_𝐵_𝑇). When a smaller energy is assumed at the pre-power stroke state, the probability of occurrence for the detached state becomes dramatically greater than the experimental result. Therefore, such binding energy seems to be necessary at the pre-power stroke state. Nevertheless, if this binding energy is not used for the force generation, it is discarded to the environment as heat. To avoid this inefficiency, we assumed the weakly binding state, from which a lever-arm swing of 30° is performed to become the pre-power stroke state. This gives one possible explanation for the Brownian search-and-catch mechanism, which assumes higher affinity to the actin filament for positive pulling forces^3, 48^.

### Statistics and reproducibility

We performed AFM experiments at least three times independently to obtain consistent results. Coordination probabilities of the AFM experiments and artificial data were statistically analyzed using Welch’s two-sided t-test. The number of independent experiments, the name of the statistical test and the exact p-values are provided in the figure legends: \**p* < 0.05, \*\**p* < 0.01, \*\*\**p* < 0.005, ****p < 0.0001.

### Data Availability

Data that support the findings of this study are available from the corresponding author upon reasonable request. All source data in the main figures are available in Source Data. Source data are provided with this paper.

## Acknowledgements

We acknowledge support by the RIKEN Quantitative Biology Center and RIKEN Junior Research Associate Program. M.I. thanks R. Kawaguchi and Y. Onishi for helping prepare the protein constructs and especially T. Ran for discussions on the muscle contraction mechanism. M.O. thanks the Bio-AFM Frontier Research Center, Kanazawa University, for technical advice about high-speed AFM. This study was supported by Grant-in-Aid for Scientific Research (B) (KAKENHI) (21H01053 to M.I.) from the Japan Society for the Promotion of Science (JSPS), AMED-PRIME from the Japan Agency for Medical Research and Development (JP19gm5810022 to M.I.), and JST, CREST Grant Number JPMJCR2023, Japan. T.W. was supported in part by the Ministry of Education, Culture, Sports, Science, and Technology of Japan (MEXT) as a Priority Issue on Post-K computing (Integrated Computational Life Science to Support Personalized and Preventive Medicine) (Project ID: hp180210 and hp190179).

## Author contributions

M.I. designed the experiments and prepared the DNA origami and protein constructs. K.I. designed the myosin constructs. H. F. and M.O. performed the experiments. H.F., K.F., H.T. and M.I. analyzed the data. T.W. carried out the numerical simulation and with M.I. analyzed the simulation data. H.F. M.O., K.F., T.W., H.T., T.Y. and M.I. interpreted the results and wrote the manuscript.

