## Supplementary information for "Dynamic coordination of the lever-arm swing of human myosin II in thick filaments on actin"

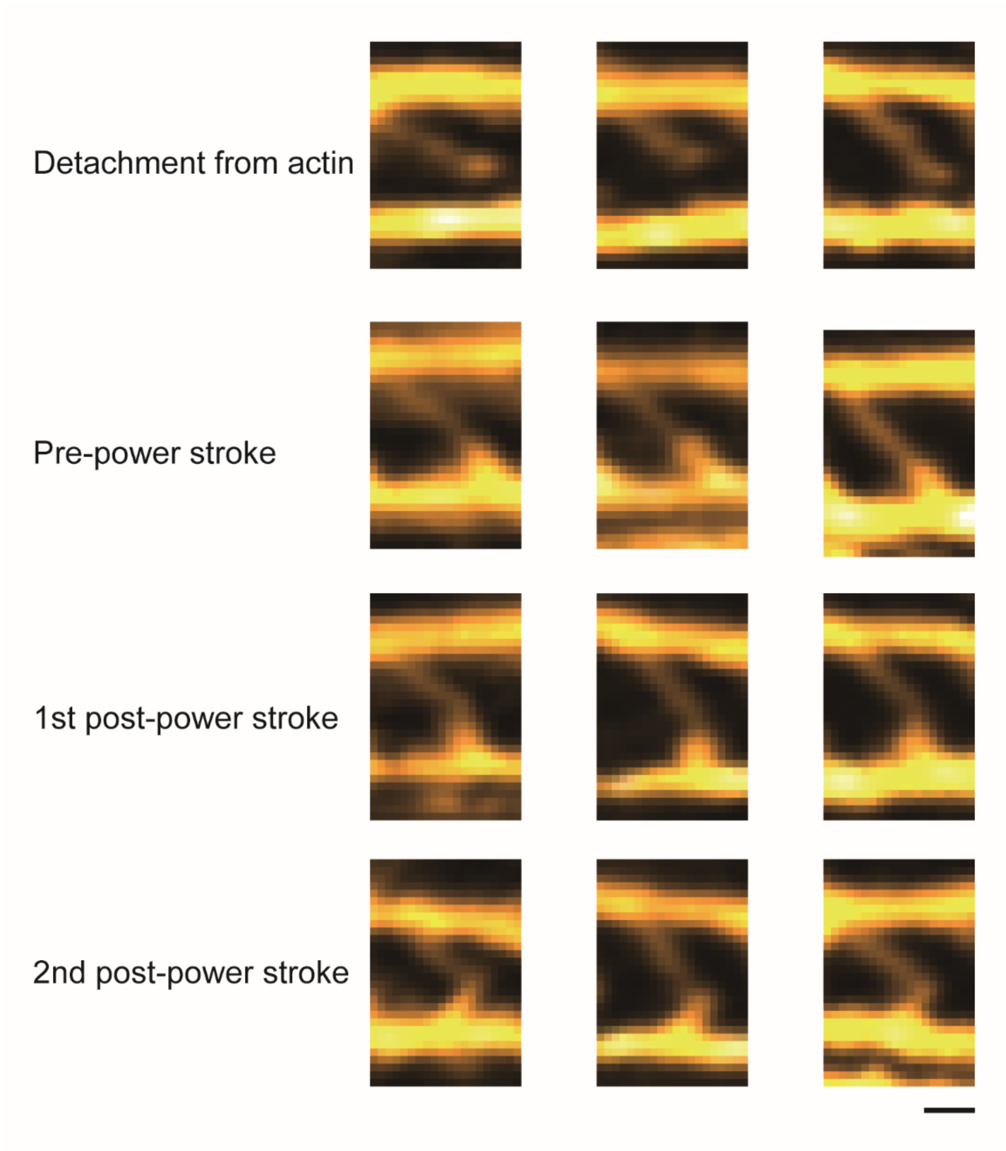


**Supplementary Figure 1. A gallery of different conformational states of myosin.** Although we did not use detailed identification for the attached state (pre, 1st and 2nd post-power stroke states), we could frequently define four states analyzed from the angle between the linker and myosin head. For the detached state, the myosin head became blurry or invisible. Scale bar, 20 nm.

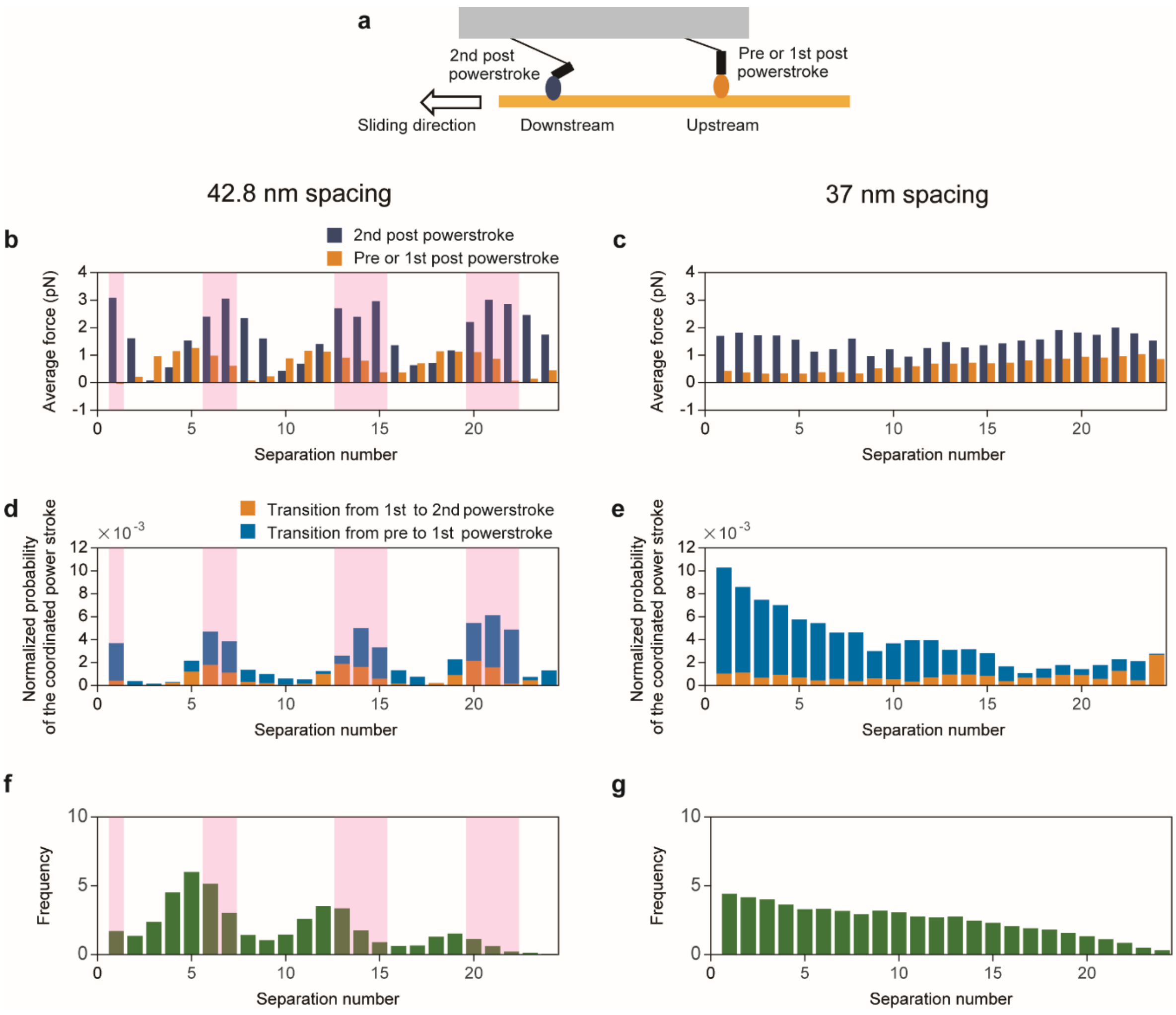


**Supplementary Figure 2. Force correlation, normalized probability of the coordinated power stroke and frequency of observed myosin pairs. a** Schematic of an analyzed myosin pair, in which the trigger myosin is downstream and the pre- or 1st post-power stroke myosin is upstream. **b and c** Average force of the trigger myosin (blue) and pre- or 1st post-power stroke myosin (orange) depending on the separation number: 42.8 nm (**b**) and 37 nm (**c**). **d and e** Normalized probability of the coordinated power stroke depending on the separation number: 42.8 nm (**d**) and 37 nm (**e**). Orange and light blue bars indicate the fraction of transitions from the pre- to 1st post-power stroke and the 1st post- to 2nd post-power stroke, respectively. **f and g** Frequency of observed myosin pairs depending on the separation number: 42.8 nm (**f**) and 37 nm (**g**). Pink areas in **b, d and f** indicate higher cooperative fraction sites (hotspots). All data were analyzed at 50 pN load.

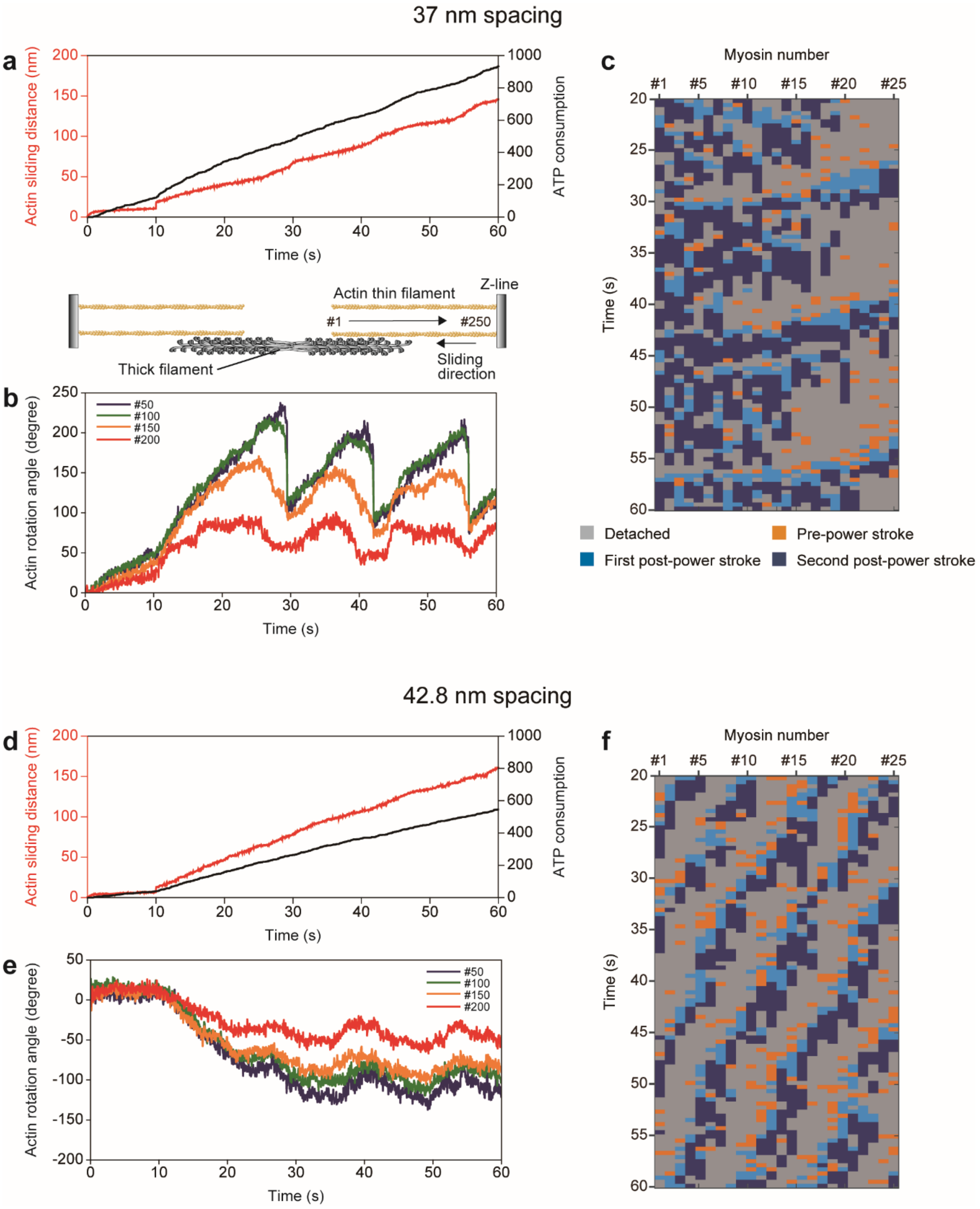
37 nm spacing
a
Actin sliding distance (nm)
ATP consumption
Time (s)
Actin thin filament
Z-line
#1
#250
Thick filament
Sliding direction
b
Actin rotation angle (degree)
#50
#100
#150
#200
Time (s)
c
Myosin number
#1
#5
#10
#15
#20
#25
Time (s)
Detached
Pre-power stroke
First post-power stroke
Second post-power stroke
42.8 nm spacing
d
Actin sliding distance (nm)
ATP consumption
Time (s)
e
Actin rotation angle (degree)
#50
#100
#150
#200
Time (s)
f
Myosin number
Time (s)

**Supplementary Figure 3. Rotation of actin filament during translocation. a-c** Time trajectory of the actin sliding distance (red), ATP consumption (black) **(a)**, rotational angle **(b)** and kymograph of the myosin molecular state **(c)** for thick filaments with 37 nm myosin spacing at 50 pN load. Colors in **b** indicate rotation of the $50^{th}$ (blue), $100^{th}$ (green), $150^{th}$ (orange) and $200^{th}$ (red) actin monomer in the actin filament (see cartoon above). The right end of the actin monomer is fixed to the Z-line; therefore, rotation at the right end is limited. Because the jump distance of myosin heads after detachment is small and the head rebinds to the same actin protofilament, actin twisting occurs and is gradually strengthened. The twist is relaxed after most myosins in the left half detach from actin. **d-f** Time trajectory of the actin sliding distance (red), ATP consumption (black) **(d)**, rotational angle **(e)** and kymograph of the myosin molecular state **(f)** for a thick filament with 42.8 nm myosin spacing at 50 pN load. Colors in **e** are the same as in **b**.

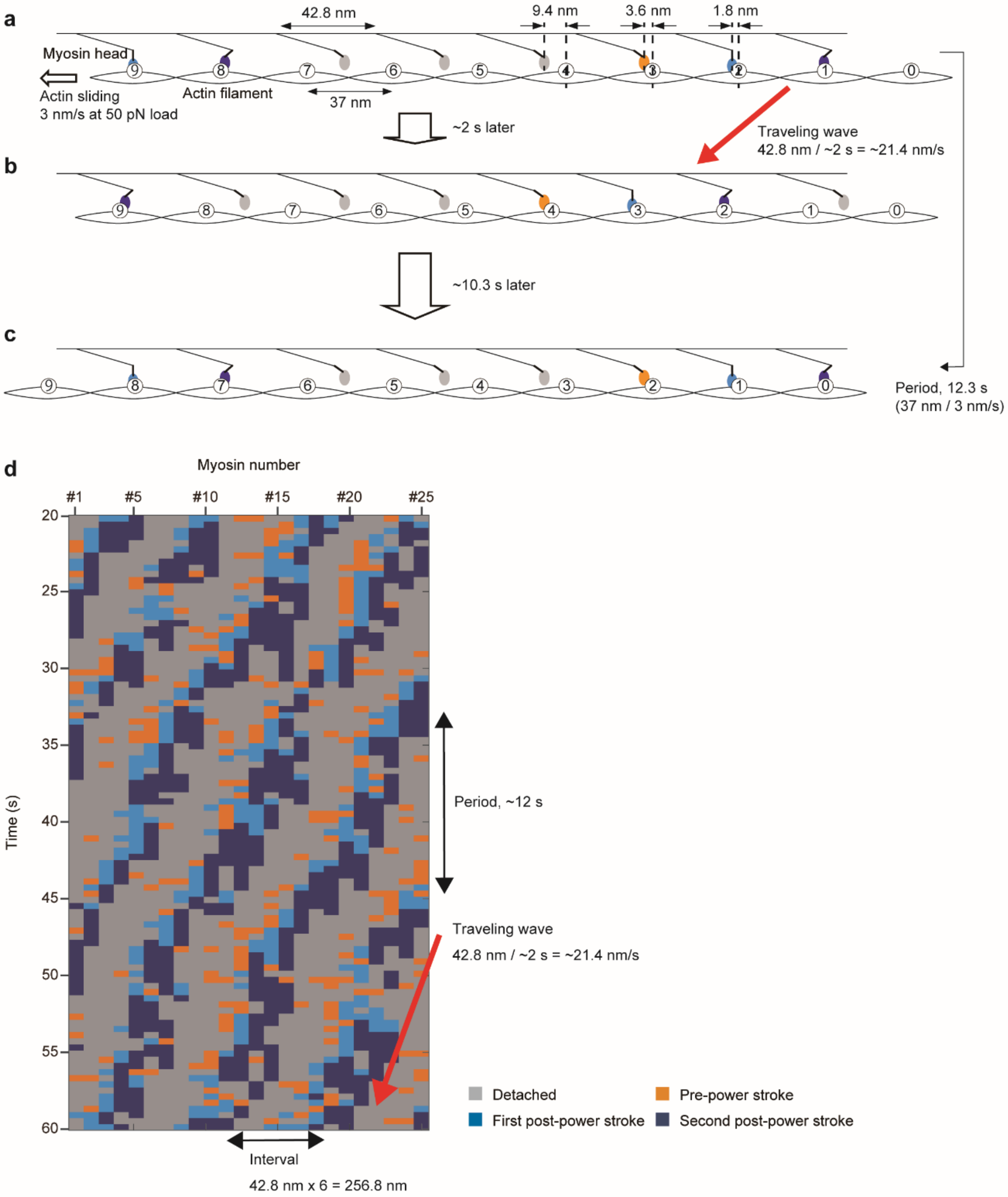
a
42.8 nm
9.4 nm
3.6 nm
1.8 nm
Myosin head
Actin sliding
3 nm/s at 50 pN load
Actin filament
37 nm
~2 s later
Traveling wave
42.8 nm / ~2 s = ~21.4 nm/s
b
~10.3 s later
c
Period, 12.3 s
(37 nm / 3 nm/s)
d
Myosin number
#1
#5
#10
#15
#20
#25
Time (s)
20
25
30
35
40
45
50
55
60
Period, ~12 s
Traveling wave
42.8 nm / ~2 s = ~21.4 nm/s
Interval
42.8 nm x 6 = 256.8 nm
Detached
Pre-power stroke
First post-power stroke
Second post-power stroke

**Supplementary Figure 4. Coordinated power strokes by thick filaments with 42.8 nm myosin spacing. a** The initial actomyosin complex. Blue myosins indicate highly accessible attached heads with actin. Due to the helical structure of the actin filament, sterically compatible binding sites are located at 37 nm intervals (numbered circles). **b** The intermediate actomyosin complex after ~2 s. **c** The actomyosin complex after 12.3 s. **d** The kymograph shown in Fig. 6d. The interval between attached states clusters, velocity of the traveling wave and period of the power stroke for a specific myosin are shown in **a-c.**

**a** Probability of three states

| | No transition | Power stroke | Detachment |
|---|---|---|---|
| Experimental data | 0.62 ± 0.08 | 0.19 ± 0.04 | 0.19 ± 0.1 |
| Artificial time series | 0.64 ± 0.01 | 0.19 ± 0.01 | 0.17 ± 0.01 |

**b** Transition fractions

Experimental data

| Pre-transition \ Post-transition | No transition | Power stroke | Detachment |
|---|---|---|---|
| No transition | 0.62 ± 0.05 | 0.24 ± 0.05 | 0.14 ± 0.06 |
| Power stroke | 0.94 ± 0.04 | 0.06 ± 0.04 | 0 |
| Detachment | 0.4 ± 0.2 | 0.1 ± 0.1 | 0.5 ± 0.3 |

Artificial time series

| Pre-transition \ Post-transition | No transition | Power stroke | Detachment |
|---|---|---|---|
| No transition | 0.62 ± 0.01 | 0.24 ± 0.01 | 0.14 ± 0.01 |
| Power stroke | 0.94 ± 0.01 | 0.06 ± 0.01 | 0 |
| Detachment | 0.38 ± 0.03 | 0.14 ± 0.02 | 0.48 ± 0.03 |

**c**

| | Number of power strokes | Number of detachments |
|---|---|---|
| Experimental data | 445 | 489 |
| Artificial time series | 466 ± 18 | 419 ± 29 |

**Supplementary Table 1. Comparison of experimental data and simulated data. a** Probability of occurrence for each state in the experiments and simulations. “No transition” is a state without a power stroke in the next frame, and “Power stroke” is a state with a power stroke in the next frame. **b** Transition fractions between states. **c** Number of transitions and power strokes produced.

### Supplementary Methods

### Three-dimensional coarse-grained model

#### A. Relationship between damping coefficient and time scale

For visual convenience, we explain the relationship between the damping coefficient and time scale by the Langevin equation with one variable:

$$\gamma \frac{dx}{dt}(t) + \frac{d\varphi}{dx}\big(x(t)\big) - R(t) = 0, \qquad \text{(S1)}$$

where $R$ is the random force that fulfills

$$\langle R(t_1)R(t_2)\rangle = 2\gamma k_B T \delta(t_1 - t_2)\ . \qquad \text{(S2)}$$

If we define the new trajectory of a particle as $\tilde{x}(t) \equiv x\big(t/\alpha_{damp}\big)$ with a time scaling constant $\alpha_{damp} > 0$, then

$$\alpha_{damp}\gamma \frac{d\tilde{x}}{dt}(t) + \frac{d\varphi}{dx}\big(\,\tilde{x}(t)\big) - R\left(\frac{t}{\alpha_{damp}}\right) = 0 \qquad \text{(S3)}$$

with

$$\langle R\left(\frac{t_1}{\alpha_{damp}}\right) R\left(\frac{t_2}{\alpha_{damp}}\right)\rangle = 2\gamma k_B T \delta\left(\frac{t_1 - t_2}{\alpha_{damp}}\right) = 2\alpha_{damp}\gamma k_B T \delta(t_1 - t_2). \qquad \text{(S4)}$$

Equation S3 with the random force given by Equation S4 is nothing but the Langevin equation with the damping coefficient $\alpha_{damp}\gamma$ for the same potential function $\varphi$ as Equation S1.

The above scaling rule on time can be understood directly from the Fokker Planck equation corresponding to Equations S1 and S2:

$$\frac{\partial}{\partial t}P(x,t) = \frac{1}{\gamma}\frac{\partial}{\partial x}\left(\frac{d\varphi}{dx}(\,x)P(x,t)\right) + \frac{k_B T}{\gamma}\frac{\partial^2}{\partial x^2}P(x,t)$$

where $P(x,t)$ is the probability of taking $x$ at time $t$. The first term on the right side corresponds to drift by the force $-\frac{d\varphi}{dx}$, and the second term corresponds to diffusion by the random force $R$.

When there are two states that switch with the rate constants $R_{ij}(x),\ 1 \le i \ne j \le 2$, the system of the Fokker Planck equations is represented by

$$\begin{cases} \frac{\partial P_1(x,t)}{\partial t} = \frac{1}{\gamma}\frac{\partial}{\partial x}\left(\frac{d\varphi_1}{dx}(\,x)P_1(x,t)\right) + \frac{k_B T}{\gamma}\frac{\partial^2 P_1(x,t)}{\partial x^2} - R_{21}(x)P_1(x,t) + R_{12}(x)P_2(x,t) \\ \frac{\partial P_2(x,t)}{\partial t} = \frac{1}{\gamma}\frac{\partial}{\partial x}\left(\frac{d\varphi_2}{dx}(\,x)P_2(x,t)\right) + \frac{k_B T}{\gamma}\frac{\partial^2 P_2(x,t)}{\partial x^2} + R_{21}(x)P_1(x,t) - R_{12}(x)P_2(x,t) \end{cases},$$

where $\varphi_i$ is the potential in the $i$-th state. If we put $\tilde{P}_i(x,t) = P_i(x, t/\alpha_{damp})$, the following equations hold.

$$\begin{cases} \frac{\partial \tilde{P}_1(x,t)}{\partial t} = \frac{1}{\tilde{\gamma}}\frac{\partial}{\partial x}\left(\frac{d\varphi_1}{dx}(\,x)\tilde{P}_1(x,t)\right) + \frac{k_B T}{\tilde{\gamma}}\frac{\partial^2 \tilde{P}_1(x,t)}{\partial x^2} - \tilde{R}_{21}(x)\tilde{P}_1(x,t) + \tilde{R}_{12}(x)\tilde{P}_2(x,t) \\ \frac{\partial \tilde{P}_2(x,t)}{\partial t} = \frac{1}{\tilde{\gamma}}\frac{\partial}{\partial x}\left(\frac{\partial d}{dx}(\,x)\tilde{P}_2(x,t)\right) + \frac{k_B T}{\tilde{\gamma}}\frac{\partial^2 \tilde{P}_2(x,t)}{\partial x^2} + \tilde{R}_{21}(x)\tilde{P}_1(x,t) - \tilde{R}_{12}(x)\tilde{P}_2(x,t) \end{cases},$$

with $\tilde{\gamma} = \alpha_{damp}\gamma$, $\tilde{R}_{ij}(x) = R_{ij}(x)/\alpha_{damp}$. These equations imply the scaling rule on time including transitions between different states.

**B. Damping coefficients**

The damping coefficients in the physiological condition were set as shown in Supplementary Table 2, where the indices of the particles are presented in Supplementary Figure 5. For the virtual particles in the myosin head, a smaller coefficient was assigned. In the HS-AFM environment, we assumed that the damping coefficient is amplified by the factor $\alpha_{damp}$=4000.

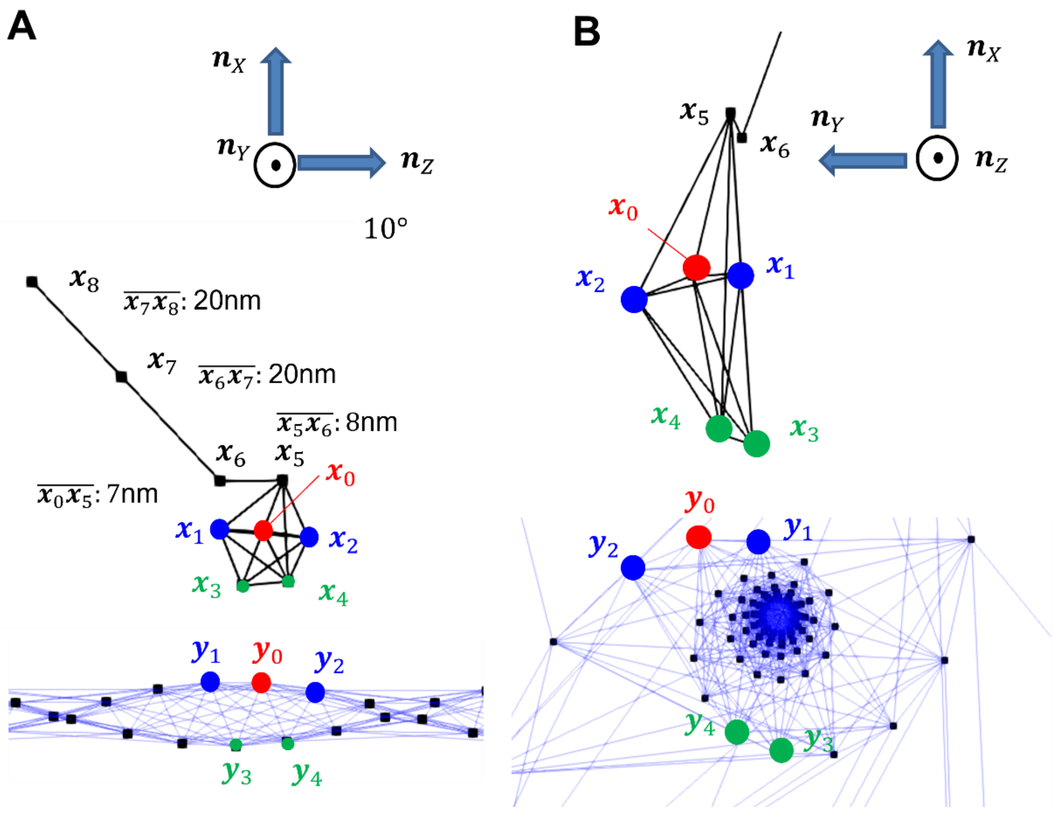


**Supplementary Figure 5.** Arrangement of particles in the myosin molecule and actin filament in the lateral view (A) and axial view (B). $\boldsymbol{n}_Z$ is the axis of the thick filament, where the myosin molecules are arranged at regular intervals. $\boldsymbol{n}_Y$ is the axis perpendicular to the lateral plane. The unloaded geometries of the myosin molecule and actin filament were defined based on the frame $\{\boldsymbol{n}_X, \boldsymbol{n}_Y, \boldsymbol{n}_Z\}$.

**Supplementary Table 2** Damping coefficients

| Particles | Damping coefficient [pN · ns/nm] |
|---|---|
| $\boldsymbol{x}_0, \boldsymbol{x}_5, \boldsymbol{x}_6, \boldsymbol{x}_7, \boldsymbol{x}_8$ | 50 |
| $\boldsymbol{x}_1, \boldsymbol{x}_2, \boldsymbol{x}_3, \boldsymbol{x}_4$ | 10 |
| $\mathbf{x}_7$ | 5 |
| $\boldsymbol{y}_0$ | 100 |

### C. Geometry

The pitch of the spiral in the actin filament was set to 36 nm without load (HS-AFM model) and 37 nm with load (sarcomere model). 13 segments connecting two adjacent actin monomers are given within one pitch. The particles that bind with the myosin heads are arranged on the cylinder with a radius of 3.5 nm (Supplementary Figure 6A). The five myosin binding particles $\boldsymbol{x}_0, \boldsymbol{x}_1, \boldsymbol{x}_2, \boldsymbol{x}_3$ and $\boldsymbol{x}_4$ are arranged so that their positions fit with the positions of the counter particles $\boldsymbol{y}_0, \boldsymbol{y}_1, \boldsymbol{y}_2, \boldsymbol{y}_3$ and $\boldsymbol{y}_4$ in the actin filament, where $\boldsymbol{y}_0$ is located at the middle point of the lateral cylindrical surface (Supplementary Figure 5). The myosin head corresponds to the line segment $\overline{\boldsymbol{x}_0\boldsymbol{x}_5}$ (length: 7 nm), which is perpendicular to $\boldsymbol{n}_Y$ and inclined at 10° from $\boldsymbol{n}_X$ (Supplementary Figure 5A). The segment $\overline{\boldsymbol{x}_5\boldsymbol{x}_6}$ (length: 8 nm) represents the lever-arm. The linker is represented by two segments $\overline{\boldsymbol{x}_6\boldsymbol{x}_7}$ (length: 20 nm) and $\overline{\boldsymbol{x}_7\boldsymbol{x}_8}$ (length: 20 nm). The root points $\boldsymbol{x}_8$ of the myosin molecules were arranged every 42.8 nm along $\boldsymbol{n}_Z$.

In the sarcomere model, the root points $\boldsymbol{x}_8$ are arranged along three edges on an equilateral triangular prism (Supplementary Figure 6B). The length of one side of the equilateral triangle is set to 40 nm. Along each edge, 25 myosin molecules are arranged every 42.8 nm, where the root point is shifted by 42.8/3 nm as it rotates the left-hand thread direction from left to right (Supplementary Figure 6C).

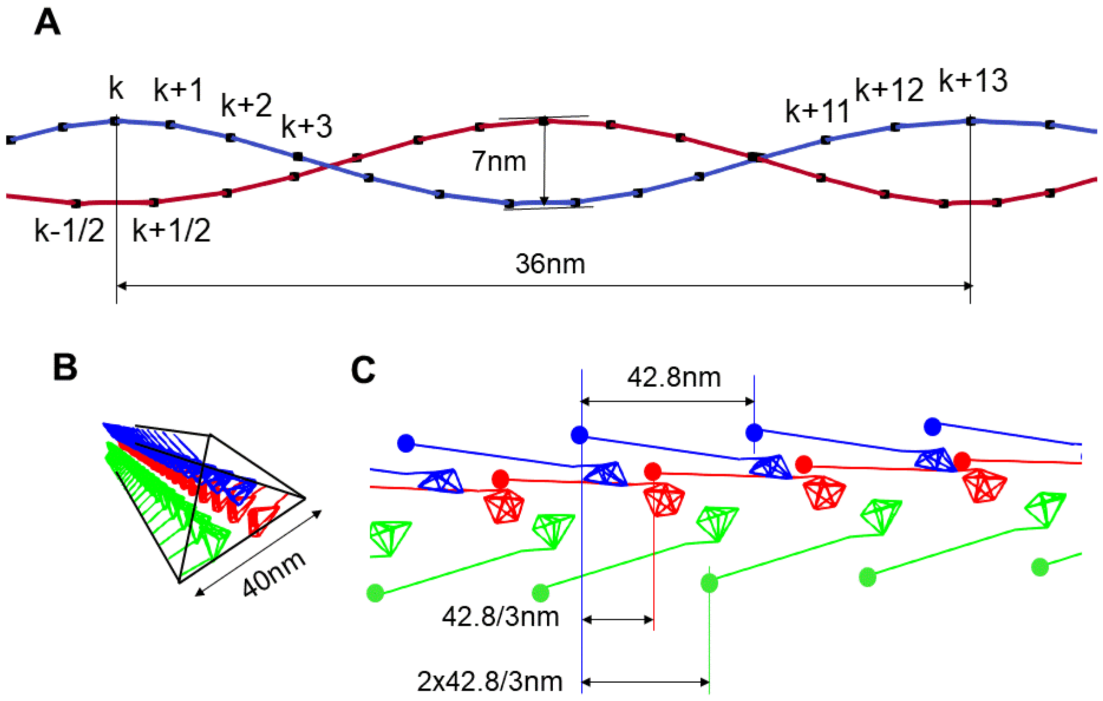


**Supplementary Figure 6** Geometry of an actin filament. Two spirals are colored blue and red, respectively. The integer indices are assigned for the binding points on the blue spiral, while the integer plus 0.5 indices are assigned for the points on the red spiral (A). The arrangement of myosin molecules in the sarcomere model (B and C).

#### D. Passive potentials of myosin molecules

In this section, the passive potential functions that maintain the myosin molecule skeletal structure are introduced. The bond length harmonic potentials are given by

$$\varphi_{M,BL,ij}(\boldsymbol{x}_i, \boldsymbol{x}_j) = \frac{1}{2} k_{M,BL,ij} \left( \|\boldsymbol{x}_i - \boldsymbol{x}_j\| - l_{ij,0} \right)^2, \qquad \text{(S5)}$$

where the natural lengths $l_{ij,0}$ are defined by the distances at the initial geometry. The spring constants $k_{M,BL,ij}$ are listed in Supplementary Table 3. For all pairs of points in the head region ($\boldsymbol{x}_0, \cdots, \boldsymbol{x}_5$), the bond length potentials are given to maintain the skeletal head structure.

**Supplementary Table 3** Coefficients of bond length potential for a myosin molecule

| Region | Index pair | $k_{M,BL,ij}$ [pN/nm] |
|---|---|---|
| Head region | $(\boldsymbol{x}_i, \boldsymbol{x}_j), 0 \le i < j \le 5$ | 50 |
| Lever-arm | $(\boldsymbol{x}_5, \boldsymbol{x}_6)$ | 50 |
| Linker | $(\boldsymbol{x}_6, \boldsymbol{x}_7)$, $(\boldsymbol{x}_7, \boldsymbol{x}_8)$ | 55 |

Some harmonic potential functions are represented as

$$\varphi_{M,Ang,X}(\theta_{M.X}) = \frac{1}{2} k_{M,Ang,X} \left( \theta_{M,X} - \theta_{M,X,0} \right)^2, \qquad \text{(S6)}$$

where $\theta_{M,X}$ is the angle defined from the relative positions of the particles in the myosin molecule. In the following, these potentials are introduced one by one, where "X" changes according to the target angle. The coefficients $k_{M,Ang,X}$ and the natural angles $\theta_{M,X,0}$ are listed in Supplementary Table 4. To represent some potentials defined by the relative position between the thick filament and the myosin molecule, we introduce the right-handed orthonormal basis $\{\boldsymbol{n}_X, \boldsymbol{n}_Y, \boldsymbol{n}_Z\}$, as shown in Supplementary Figure 7. Here, $\boldsymbol{n}_Z$ represents the unit vector that directs the thick filament from left to right. The plane spanned by $\boldsymbol{n}_X$ and $\boldsymbol{n}_Z$ determines the position of the linker particles in the unloaded condition. Note that $\boldsymbol{n}_X$ and $\boldsymbol{n}_Y$ must be defined separately for each thick filament in the case of the sarcomere model.

[**Angle between thick filament backbone and linker** ($\theta_{M,TFLI}$)]

This potential represents the stiffness associated with the angle between the thick filament backbone and the linker (Supplementary Figure 7A). Thus, it is given based on the unit vector $\boldsymbol{d} = \frac{\boldsymbol{x}_7 - \boldsymbol{x}_8}{\|\boldsymbol{x}_7 - \boldsymbol{x}_8\|}$. Basically, $\theta_{M,TFLI}$ satisfies the following relationship.

$$B \coloneqq \boldsymbol{d} \cdot \boldsymbol{n}_Z = \cos\theta_{M,TFLI}. \qquad \text{(S7)}$$

Because the sign of $\theta_{M,TFLI}$ is indefinite with Equation S7, we also refer to the following relationship.

$$A := \sin\theta_{M,TFLI} = \begin{cases} \sqrt{1 - B^2}, \boldsymbol{d} \cdot \boldsymbol{n}_X \leq 0 \\ -\sqrt{1 - B^2}, \boldsymbol{d} \cdot \boldsymbol{n}_X > 0 \end{cases}. \qquad \text{(S8)}$$

Using the variables $A$ and $B$, $\theta_{M,TFLI}$ is calculated as follows.

$$\theta_{M,TFLI} = \begin{cases} \arcsin A\,, \boldsymbol{d} \cdot \boldsymbol{n}_Z > 0 \quad (-\pi/2 < \theta_{M,TFLI} < \pi/2) \\ \arccos B\,, \boldsymbol{d} \cdot \boldsymbol{n}_Z \leq 0, \boldsymbol{d} \cdot \boldsymbol{n}_X \leq 0 \ \ (\pi/2 \leq \theta_{M,TFLI} \leq \pi) \\ -\arccos B\,, \boldsymbol{d} \cdot \boldsymbol{n}_Z \leq 0, \boldsymbol{d} \cdot \boldsymbol{n}_X > 0 \ \ (-\pi < \theta_{M,TFLI} \leq -\pi/2) \end{cases}. \qquad \text{(S9)}$$

As shown in Supplementary Table 4, the different coefficients were adopted for $\theta_{M,TFLI} \leq \theta_{M,TFLI,0}$ and $\theta_{M,TFLI} > \theta_{M,TFLI,0}$ to reproduce the distribution obtained by Fujita et al.[1].

[**Angle between XZ-plane and linker** ($\theta_{M,XZLI}$)]

This potential keeps the linker staying in the plane spanned by $\boldsymbol{n}_X$ and $\boldsymbol{n}_Z$ (Supplementary Figure 7A). The angle $\theta_{M,XZLI}$ is defined by

$$\theta_{M,XZLI} = \arcsin \boldsymbol{d} \cdot \boldsymbol{n}_Y. \qquad \text{(S10)}$$

[**Bending of linker** ($\theta_{M,BELI}$)]
This potential gives the stiffness for bending of the linker. The angle $\theta_{M,BELI}$ is defined by the two direction vectors, $\boldsymbol{d} = \frac{\boldsymbol{x}_7 - \boldsymbol{x}_8}{\|\boldsymbol{x}_7 - \boldsymbol{x}_8\|}$ and $\boldsymbol{e} = \frac{\boldsymbol{x}_6 - \boldsymbol{x}_7}{\|\boldsymbol{x}_6 - \boldsymbol{x}_7\|}$, as follows (Supplementary Figure 7A).

$$\theta_{M,BELI} = \arccos \boldsymbol{d} \cdot \boldsymbol{e}. \qquad \text{(S11)}$$

[**Angle between linker and lever-arm** ($\theta_{M,LILA}$)]
The angle $\theta_{M,LILA}$ is defined by projecting the direction vector of the linker ($\boldsymbol{e} = \frac{\boldsymbol{x}_6 - \boldsymbol{x}_7}{\|\boldsymbol{x}_6 - \boldsymbol{x}_7\|}$) and the direction vector of the lever-arm ($\boldsymbol{f} = \frac{\boldsymbol{x}_5 - \boldsymbol{x}_6}{\|\boldsymbol{x}_5 - \boldsymbol{x}_6\|}$) to the plane determined by the three points $\boldsymbol{x}_1$, $\boldsymbol{x}_2$, and $\boldsymbol{x}_5$ contained in the head region (Supplementary Figure 7B).

$$\theta_{M,LILA} = \begin{cases} \arcsin(\boldsymbol{w} \cdot \boldsymbol{e} \times \boldsymbol{f}), & \boldsymbol{e} \cdot \boldsymbol{f} > 0 \ \ (-\pi/2 < \theta_{M,LILA} < \pi/2) \\ \pi - \arcsin(\boldsymbol{w} \cdot \boldsymbol{e} \times \boldsymbol{f}), & \boldsymbol{e} \cdot \boldsymbol{f} \le 0 \ (\pi/2 \le \theta_{M,LILA} < 3\pi/2) \end{cases}. \qquad \text{(S12)}$$

Here, the direction vector $\boldsymbol{w}$ is given by

$$\boldsymbol{w} = \frac{(\boldsymbol{x}_5 - \boldsymbol{x}_1) \times (\boldsymbol{x}_2 - \boldsymbol{x}_1)}{\|(\boldsymbol{x}_5 - \boldsymbol{x}_1) \times (\boldsymbol{x}_2 - \boldsymbol{x}_1)\|}. \qquad \text{(S13)}$$

Note that the angle $\theta_{M,LILA}$ is same as the angle $\theta$, which is used to represent the distribution of the attached myosin molecules.

[**Constraints of the lever-arm** ($\theta_{M,CLA}$)]
The angle $\theta_{M,CLA}$ represents the deviation of the lever-arm from the plane determined by $\boldsymbol{x}_1$, $\boldsymbol{x}_2$, and $\boldsymbol{x}_5$ contained in the head region (Supplementary Figure 7B).

$$\theta_{M,CLA} = \arcsin \boldsymbol{f} \cdot \boldsymbol{w}. \qquad \text{(S14)}$$

To avoid extraordinary lever-arm rotation with respect to the myosin head, a constraint is imposed on the angle $\theta_{M,HLA}$ defined by

$$\theta_{M,HLA} = \arcsin \boldsymbol{f} \cdot \boldsymbol{h}, \qquad \text{(S15)}$$

where $\boldsymbol{h} = \frac{\boldsymbol{x}_0 - \boldsymbol{x}_5}{\|\boldsymbol{x}_0 - \boldsymbol{x}_5\|}$ is the direction vector of the myosin head (Supplementary Figure 7B).

The constraint is imposed only for $\theta_{M,HLA} < \theta_{M,HLA,0}$ by the potential:

$$\varphi_{M,Ang,HLA}(\theta_{M.HAL}) = \begin{cases} \frac{1}{2} k_{M,Ang,HLA}\left(\theta_{M,HLA} - \theta_{M,HLA,0}\right)^2, \theta_{M,HLA} < \theta_{M,HLA,0}, \\ 0, \quad \theta_{M,HLA} \geq \theta_{M,HLA,0} \end{cases}. \qquad \text{(S16)}$$

[**Flapping and twisting of a myosin head** ($\theta_{M,FLAP}$ and $\theta_{M,TWST}$)]

To repress the flapping of a myosin head at the junction with the linker (Supplementary Figure 7B), we introduce stiffness associated with the flapping angle:

$$\theta_{M,FLAP} = \arcsin \boldsymbol{e} \cdot \boldsymbol{v}, \qquad \text{(S17)}$$

where the unit vector $\boldsymbol{v}$ is given by

$$\boldsymbol{v} = \frac{(\boldsymbol{x}_6 - \boldsymbol{x}_1) \times (\boldsymbol{x}_2 - \boldsymbol{x}_1)}{\|(\boldsymbol{x}_6 - \boldsymbol{x}_1) \times (\boldsymbol{x}_2 - \boldsymbol{x}_1)\|}. \qquad \text{(S18)}$$

Similarly, to repress twisting of the myosin head at the junction with the linker, we introduce stiffness associated with the twisting angle:

$$\theta_{M,TWST} = \arcsin \boldsymbol{g} \cdot \boldsymbol{v}, \qquad \text{(S19)}$$

where the unit vector $\boldsymbol{g}$ is given by

$$\boldsymbol{g} = \frac{(\boldsymbol{x}_6 - \boldsymbol{x}_8) \times \boldsymbol{n}_Y}{\|(\boldsymbol{x}_6 - \boldsymbol{x}_8) \times \boldsymbol{n}_Y\|}. \qquad \text{(S20)}$$

[**Rigorous definition of power stroke rotation angle** $\eta$]

Here, the rigorous definition of the power stroke rotation angle $\eta$ is given. This variable is used to determine the active potentials $\varphi_{PS}$ and $\varphi_{RS}$. $\eta$ is defined so that it satisfies

$$\sin\eta = \boldsymbol{l}\cdot\boldsymbol{n}, \quad \text{(S21)}$$

with the two direction vectors $\boldsymbol{n} = \frac{\boldsymbol{x}_2-\boldsymbol{x}_1}{\|\boldsymbol{x}_2-\boldsymbol{x}_1\|}$ and $\boldsymbol{l} = \frac{\boldsymbol{x}_6-\boldsymbol{x}_5}{\|\boldsymbol{x}_6-\boldsymbol{x}_5\|}$. As shown in Supplementary Figure 7C, to give a rigorous definition of $\eta$, we use the direction vector $\boldsymbol{m}$ perpendicular to $\boldsymbol{n}$. Namely, $\boldsymbol{m}$ is constructed as

$$\boldsymbol{m} = \frac{\boldsymbol{t}-(\boldsymbol{t}\cdot\boldsymbol{n})\boldsymbol{n}}{\|\boldsymbol{t}-(\boldsymbol{t}\cdot\boldsymbol{n})\boldsymbol{n}\|} \quad \text{(S22)}$$

with $\boldsymbol{t} = \boldsymbol{x}_5 - \frac{1}{2}(\boldsymbol{x}_1+\boldsymbol{x}_2)$. Then, the rotation angle $\eta$ is defined by

$$\eta = \begin{cases} \arcsin\boldsymbol{l}\cdot\boldsymbol{n},\ \ \boldsymbol{l}\cdot\boldsymbol{m}\geq 0 \quad (-\pi/2\leq\eta\leq\pi/2) \\ -\pi-\arcsin\boldsymbol{l}\cdot\boldsymbol{n}, \boldsymbol{l}\cdot\boldsymbol{m}<0, \boldsymbol{l}\cdot\boldsymbol{n}<0(-\pi<\eta<-\pi/2). \\ \pi-\arcsin\boldsymbol{l}\cdot\boldsymbol{n}, \boldsymbol{l}\cdot\boldsymbol{m}<0, \boldsymbol{l}\cdot\boldsymbol{n}\geq 0(\pi/2<\eta\leq\pi) \end{cases} \quad \text{(S23)}$$

**Supplementary Table 4** Material parameters for the angle stiffness of a myosin molecule

| Coefficient name | Related points | Coefficient [pN nm/rad] | Natural angle [deg] |
|---|---|---|---|
| $k_{M,TFLI}$ | $\boldsymbol{x}_7, \boldsymbol{x}_8$ | 160 ($\theta_{M,TFLI} \leq \theta_{M,TFLI,0}$)<br>320 ($\theta_{M,TFLI} > \theta_{M,TFLI,0}$) | 47 (HS-AFM),<br>15 (Sarcomere) |
| $k_{M,XZLI}$ | $\boldsymbol{x}_7, \boldsymbol{x}_8$ | 400 | 0 |
| $K_{M,BELI}$ | $\boldsymbol{x}_6, \boldsymbol{x}_7, \boldsymbol{x}_8$ | 100 | 0 |
| $k_{M,LILA}$ | $\boldsymbol{x}_1, \boldsymbol{x}_2, \boldsymbol{x}_5, \boldsymbol{x}_6, \boldsymbol{x}_7$ | 10 | 0 (HS-AFM)<br>60 (Sarcomere) |
| $K_{M,CLA}$ | $\boldsymbol{x}_1, \boldsymbol{x}_2, \boldsymbol{x}_5, \boldsymbol{x}_6$ | 1000 | 0 |
| $K_{M,HLA}$ | $\boldsymbol{x}_0, \boldsymbol{x}_5, \boldsymbol{x}_6$ | 1000 | –10 |
| $K_{M,FLAP}$ | $\boldsymbol{x}_1, \boldsymbol{x}_2, \boldsymbol{x}_6, \boldsymbol{x}_7$ | 400 | 0 |
| $K_{M,TWST}$ | $\boldsymbol{x}_1, \boldsymbol{x}_2, \boldsymbol{x}_6, \boldsymbol{x}_8$ | 400 | 0 |

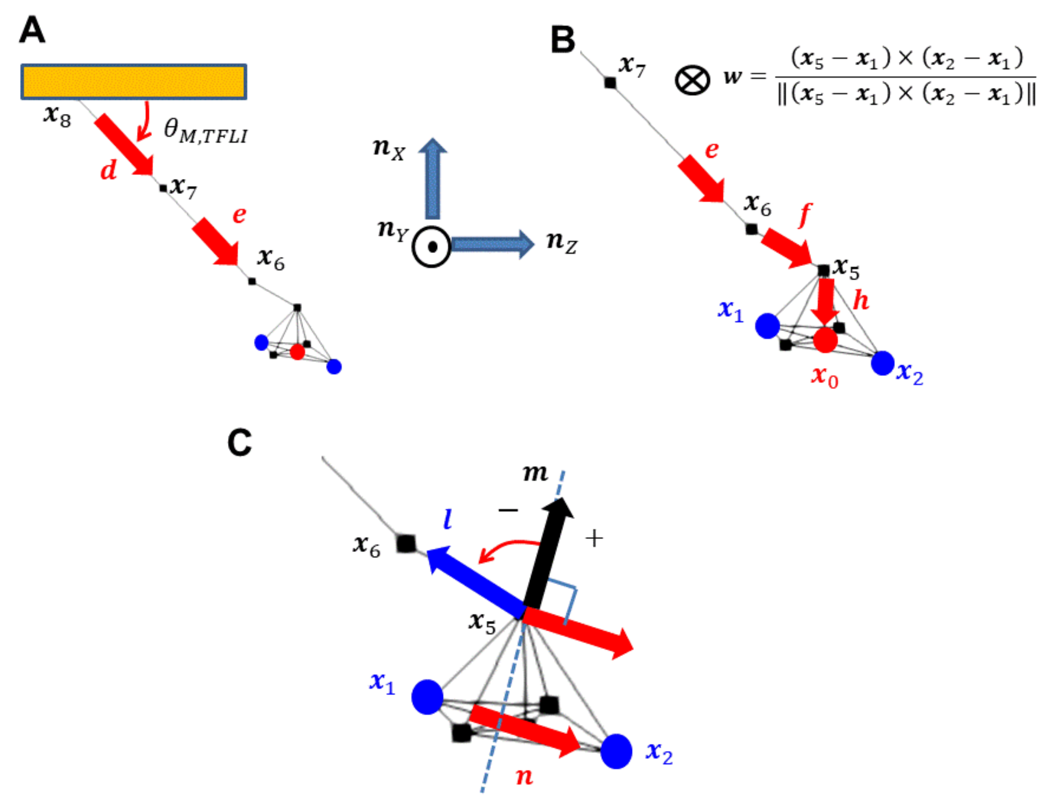


**Supplementary Figure 7** Direction vectors to define the stiffness of a myosin molecule. The unit vector $\boldsymbol{d}$ is used to define the angle $\theta_{M,TFLI}$ between the thick filament backbone and linker and the angle $\theta_{M,XZLI}$ between the XZ-plane and linker. The unit vectors $\boldsymbol{d}$ and $\boldsymbol{e}$ are used to define the bending angle $\theta_{M,BELI}$ of the linker (A). The unit vectors $\boldsymbol{e}$, $\boldsymbol{f}$, and $\boldsymbol{w}$ are used to define the angle $\theta_{M,LILA}$ between the linker and lever-arm. The unit vectors $\boldsymbol{f}$ and $\boldsymbol{h}$ are used to define the angle $\theta_{M,HLA}$ between the lever-arm and myosin head. The angle $\theta_{M,HLA}$ is used to avoid extraordinary lever-arm rotation (B). The unit vectors $\boldsymbol{l}$, **m**, and $\boldsymbol{n}$ are used to define the power stroke rotation angle $\eta$ (C).

### E. Stiffness of an actin filament

Bond length harmonic potentials with the coefficient 60 pN/nm were imposed as with Equation S5 for the pairs (*k*,*k*+1) and (*k*,*k*+3) in the same spiral, and for the pairs (*k*,*k*+1/2), (*k*,*k*+3/2), (*k*,*k*+5/2), and (*k*,*k*+7/2) between two spirals with respect to the indices in Supplementary Figure 8. In addition, the dihedral angle harmonic potentials with the coefficient 2000 pN nm/rad were imposed for the combinations (*k*,*k*+1/2,*k*+1,*k*+3/2) and (*k*-

1,*k*+1/2,*k*+2,*k*+7/2). The material parameters were determined so that the actual persistence length of an actin filament (15 μm)[2] was reproduced.

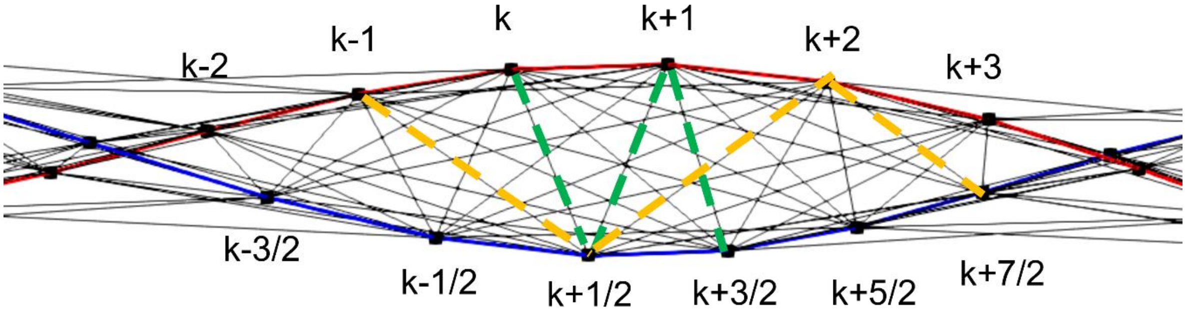


**Supplementary Figure 8** Bonds (black lines) and the dihedral angle combinations (green and yellow broken lines) in the actin filament. The binding points are indexed with integers on one spiral and with integers +1/2 on the other spiral.

### F. Exclusive potential between a myosin and actin filament

Between points, except for the virtual points $\boldsymbol{x}_1, \boldsymbol{x}_2, \boldsymbol{x}_3$ and $\boldsymbol{x}_4$, in the myosin molecule and points in the actin filament, exclusive potentials are imposed. The following cylindrical exclusive potential associated with point $\boldsymbol{y}_0$ is superposed over the actin filament (Supplementary Figure 9). The central point of the cylinder is defined by

$$\boldsymbol{y}_c = \frac{1}{2}\boldsymbol{y}_0 + \frac{1}{4}(\boldsymbol{y}_3 + \boldsymbol{y}_4). \quad \text{(S24)}$$

The longitudinal direction vector is defined by

$$\boldsymbol{a} = \frac{\boldsymbol{y}_2 - \boldsymbol{y}_1}{\|\boldsymbol{y}_2 - \boldsymbol{y}_1\|}. \quad \text{(S25)}$$

Let us define the radial and longitudinal coordinates for $\boldsymbol{x}$ by

$$\begin{cases} r(\boldsymbol{x}, \boldsymbol{Y}) = \|\boldsymbol{x} - \boldsymbol{y}_c - \{(\boldsymbol{x} - \boldsymbol{y}_c) \cdot \boldsymbol{a}\}\boldsymbol{a}\| \\ s(\boldsymbol{x}, \boldsymbol{Y}) = (\boldsymbol{x} - \boldsymbol{y}_c) \cdot \boldsymbol{a} \end{cases}, \quad \text{(S26)}$$

with $\boldsymbol{Y} = [\boldsymbol{y}_0, \boldsymbol{y}_1, \boldsymbol{y}_2, \boldsymbol{y}_3, \boldsymbol{y}_4]$.

Our exclusive potential is given as follows,

$$\varphi_C(\boldsymbol{x}, \boldsymbol{Y}) = F_{barrier}\big(r(\boldsymbol{x}, \boldsymbol{Y})\big)G_{barrier}\big(s(\boldsymbol{x}, \boldsymbol{Y})\big) \quad \text{(S27)}$$

with

$$F_{barrier}(r) = \begin{cases} \frac{1}{2}k_r(r - R_0)^2, r < R_0 \\ 0, r \geq R_0 \end{cases} \quad \text{(S28)}$$

$$G_{barrier}(s) = \begin{cases} 0, s \leq -S_0 \\ \frac{1}{2}k_s(s + S_0)^2, -S_0 < s \leq 0 \\ \frac{1}{2}k_s(s - S_0)^2, 0 < s \leq S_0 \\ 0, s > S_0 \end{cases}, \quad \text{(S29)}$$

where $R_0$ =3 nm, $S_0$=5.5 nm, $k_r$=100 pN/nm, and $k_s$=50 pN/nm adopted in the numerical simulations.

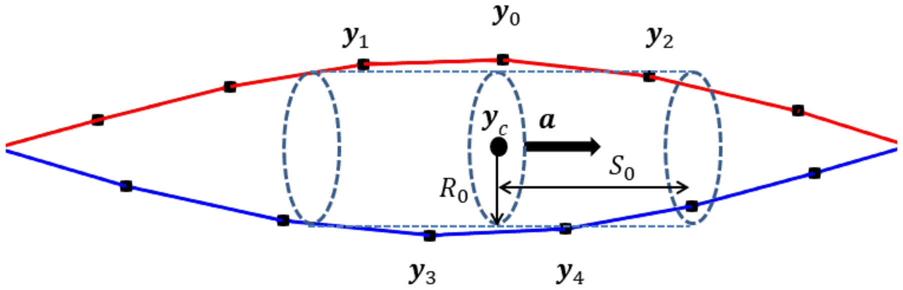


**Supplementary Figure 9** Cylindrical region of the exclusive potential for the binding point $\boldsymbol{y}_0$. The cylinder represents the potential barrier that inhibits penetration of the points $\boldsymbol{x}_0$, $\boldsymbol{x}_5$, and $\boldsymbol{x}_6$ in the myosin molecule into the thin filament.

### G. Power stroke and recovery stroke active potential functions

The power stroke and recovery stroke potentials $\varphi_{PS}$ and $\varphi_{RS}$ are given as the function of two variables $(\eta, \chi)$, where $\eta$ is the lever-arm rotation angle as defined in Equation S23 and $\chi$ is the binding degree (Equation 5 in the main text). The potentials are given by combining the local minimal energy functions associated with different states. The parameters to define the local minimal energy functions are listed in Supplementary Table 5. The basic part of the power stroke potential $\varphi_{PS}$ is defined by combining the following five local minimal energy functions:

$$\begin{cases} \varphi_{PS,Det}(\eta,\chi) = E_{Det}\lambda_{A_{Det},B_{Det},\eta_{Det}}(\eta)\nu_{C_{Det}}(\chi) \\ \varphi_{PS,WB}(\eta,\chi) = E_{WB}\lambda_{A_{WB},B_{WB},\eta_{WB}}(\eta)\mu_{C_{WB},\chi_{WB},}(\chi) \\ \varphi_{PS,Pre}(\eta,\chi) = E_{Pre}\lambda_{A_{Pre},B_{Pre},\eta_{Pre}}(\eta)\mu_{C_{Pre},\chi_{Pre}}(\chi) \\ \varphi_{PS,1st}(\eta,\chi) = E_{1st}\lambda_{A_{1st},B_{1st},\eta_{1st}}(\eta)\mu_{C_{1st},\chi_{1st}}(\chi) \\ \varphi_{PS,2nd}(\eta,\chi) = E_{2nd}\lambda_{A_{2nd},B_{2nd},\eta_{2nd}}(\eta)\mu_{C_{1st},\chi_{2nd}}(\chi) \end{cases}, \qquad \text{(S30)}$$

where $E_{Det}$, $E_{WB}$, $E_{Pre}$, $E_{1st}$ and $E_{2nd}$ denote the local minimal energies in the force producing phase, respectively, at the detached state, the weakly binding state, the pre-power stroke state, the first post-power stroke state, and the second post-power stroke state. The three functions $\lambda_{A,B,\eta_0}(\eta)$ , $\nu_C(\chi)$ and $\mu_{C,\chi_0}(\chi)$ are given along the $\eta$- and $\chi$-coordinates, respectively, by

$$\lambda_{A,B,\eta_0}(\eta) = \begin{cases} 0, \eta < \eta_0 - A \\ (1+\cos(\pi(\eta-\eta_0)/A))/2\,, \eta_0 - A \le \eta < \eta_0 \\ (1+\cos(\pi(\eta-\eta_0)/B))/2\,, \eta_0 \le \eta < \eta_0 + B \\ 0, \eta \ge \eta_0 + B \end{cases} \qquad \text{(S31)}$$

$$\nu_C(\chi) = \begin{cases} 1, \chi < 0 \\ (1+\cos(\pi\chi/C))/2\,, 0 \le \chi < C \\ 0, \chi \ge C \end{cases} \qquad \text{(S32)}$$

$$\mu_{C,\chi_0}(\chi) = \begin{cases} 0, \chi < \chi_0 - C \\ (1+\cos(\pi(\chi-\chi_0)/C))/2\,, \chi_0 - C \le \chi < \chi_0. \\ 1, \chi \ge \chi_0 \end{cases} \qquad \text{(S33)}$$

The product of the two functions, $\lambda_{A,B,\eta_0}(\eta)$ and $\mu_{C,\chi_0}(\chi)$, in Equation S30 have extreme values at $(\eta_0,\chi)$, $\chi \ge \chi_0$. The parameters $A$, $B$, and $C$ represent the reciprocal curvatures of the potential around the extremes.

The integration of the five functions in Equation S30 is performed based on combining the two functions $\varphi_1$ and $\varphi_2$.

$$\Psi_D(\varphi_1, \varphi_2) = \frac{1}{2}\left(\varphi_1 + \varphi_2 - \sqrt{(\varphi_1 - \varphi_2)^2 + 4D^2}\right) + D. \qquad \text{(S34)}$$

Here, the parameter $D > 0$ has the dimension of energy and corresponds to the difference between the junction value $\varphi_1 = \varphi_2$ and $\Psi_D(\varphi_1, \varphi_2)$ at the barrier that separates the two local minimums of potentials $\varphi_1$ and $\varphi_2$. The last term on the right side of Equation S34 is added to make $\Psi_D$ equal to zero on the region where both $\varphi_1$ and $\varphi_2$ are zero. To combine the five functions, first, two pairs $\{\varphi_{PS,WB}, \varphi_{PS,Pre}\}$ and $\{\varphi_{PS,1st}, \varphi_{PS,2nd}\}$ are combined as

$$\begin{cases} \Psi_{WB,Pre}(\eta,\chi) = \Psi_{D_{WB,Pre}}\left(\varphi_{PS,WB}(\eta,\chi), \varphi_{PS,Pre}(\eta,\chi)\right) \\ \Psi_{1st,2nd}(\eta,\chi) = \Psi_{D_{1st,2nd}}\left(\varphi_{PS,1st}(\eta,\chi), \varphi_{PS,2nd}(\eta,\chi)\right) \end{cases}. \qquad \text{(S35)}$$

The first function in Equation S35 is further combined with $\varphi_{PS,Det}$ as

$$\Psi_{Det,WB,Pre}(\eta,\chi) = \Psi_{D_{Det,WB,Pre}}\left(\varphi_{PS,Det}(\eta,\chi), \Psi_{WB,Pre}(\eta,\chi)\right). \qquad \text{(S36)}$$

Finally, the five functions are combined as

$$\Psi_{PS}(\eta,\chi) = \Psi_{D_{PS}}\left(\Psi_{Det,WB,Pre}(\eta,\chi), \Psi_{1st,2nd}(\eta,\chi)\right). \qquad \text{(S33)}$$

Though $\Psi_{PS}$ has the capability to represent the landscape of the potential at the binding states, the flat landscape for $\chi \approx 0$ does not reflect the behavior of a myosin molecule in the detached state, where the lever-arm rotation angle is likely preserved. In our numerical model, we add barriers to prevent the lever-arm rotation in the detachedstate. The local minimums of the detached states are given by

$$\begin{cases} \alpha_{Pre}(\eta,\chi) = E_{DPre}\lambda_{A_{DPre},B_{DPre}}(\eta-\eta_{DPre})\nu_{C_{DPre}}(\chi) \\ \alpha_{1st}(\eta,\chi) = E_{D1st}\lambda_{A_{D1st},B_{D1st}}(\eta-\eta_{D1st})\nu_{C_{D1st}}(\chi) \\ \alpha_{2nd}(\eta,\chi) = E_{D2nd}\lambda_{A_{D2nd},B_{D2nd}}(\eta-\eta_{D2nd})\nu_{C_{D2nd}}(\chi) \end{cases}. \qquad \text{(S34)}$$

These three local minimums in the detached state and the local minimum defined by $\varphi_{PS,Det}(\eta,\chi)$ in Equation S30 are separated by the following three barriers.

$$\begin{cases} \beta_{Pre}(\eta,\chi) = E_{HPre}\lambda_{A_{HPre},B_{HPre}}(\eta-\eta_{HPre})\nu_{C_{HPre}}(\chi) \\ \beta_{1st}(\eta,\chi) = E_{H1st}\lambda_{A_{H1st},B_{H1st}}(\eta-\eta_{H1st})\nu_{C_{H1st}}(\chi) \\ \beta_{2nd}(\eta,\chi) = E_{H2nd}\lambda_{A_{H2nd},B_{H2nd}}(\eta-\eta_{H2nd})\nu_{C_{H2nd}}(\chi) \end{cases}. \qquad \text{(S35)}$$

Finally, the active potential of the power stoke is defined by

$$\varphi_{PS}(\eta,\chi) = \Psi_{PS}(\eta,\chi) + \alpha_{Pre}(\eta,\chi) + \alpha_{1st}(\eta,\chi) + \alpha_{2nd}(\eta,\chi) + \beta_{Pre}(\eta,\chi) + \beta_{1st}(\eta,\chi) + \beta_{2nd}(\eta,\chi). \qquad \text{(S36)}$$

The active potential of the recovery power stroke is defined by adding two functions,

$$\varphi_{RS}(\eta,\chi) = \alpha_{RS,\mathrm{RD}}(\eta,\chi) + \alpha_{RS,\mathrm{R2}}(\eta,\chi), \qquad \text{(S37)}$$

where the two functions are given by

$$\begin{cases} \alpha_{RS,RD}(\eta,\chi) = E_{RD}\lambda_{A_{RD},B_{RD},\eta_{RD}}(\eta)\nu_{C_{RD}}(\chi) \\ \alpha_{RS,R2}(\eta,\chi) = E_{R2}\lambda_{A_{R2},B_{R2},\eta_{R2}}(\eta)\mu_{C_{R2},\chi_{R2},}(\chi) \end{cases}. \qquad \text{(S38)}$$

**Supplementary Table 5** Parameters to define the power stroke potential $\varphi_{PS}$ and the recovery power stroke potential $\varphi_{RS}$. Note the energy parameters $E_*$ are not equal to the local minimums of $\varphi_{PS}$.

**Power stroke potential**

Local minimum at attachment (Equation S30).

| **Function** | $E_*$ $[k_BT]$ | $A_*$ [°] | $B_*$ [°] | $\eta_*$ [°] | $C_*$ | $\chi_*$ |
|---|---|---|---|---|---|---|
| $\varphi_{PS,Det}$ | –4 | 24 | 24 | –90 | 0.4 | |
| $\varphi_{PS,WB}$ | –8 | 24 | 24 | –90 | 0.8 | 0.45 |

| | | | | | | |
|---|---|---|---|---|---|---|
| $\varphi_{PS,Pre}$ | −15 | 34 | 48 | −60 | 0.8 | 0.6 |
| $\varphi_{PS,1st}$ | −18.5 | 50 | 34 | 0 | 0.8 | 0.85 |
| $\varphi_{PS,2nd}$ | −29 | 34 | 34 | 40 | 0.8 | 0.9 |

Local minimum at detachment (Equation S34).

| **Function** | $E_{D*}$ $[k_B T]$ | $A_{D*}$ [°] | $B_{D*}$ [°] | $\eta_{D*}$ | $C_{D*}$ |
|---|---|---|---|---|---|
| $\alpha_{Pre}$ | −3 | 20 | 20 | −60 | 0.1 |
| $\alpha_{1st}$ | −3 | 20 | 20 | 0 | 0.1 |
| $\alpha_{2nd}$ | −3 | 20 | 20 | 40 | 0.1 |

Barrier function at detachment (Equation S35).

| **Function** | $E_{H*}$ $[k_B T]$ | $A_{H*}$ [°] | $B_{H*}$ [°] | $\eta_{H*}$ | $C_{H*}$ |
|---|---|---|---|---|---|
| $\beta_{Pre}$ | 30 | 20 | 20 | −75 | 0.6 |
| $\beta_{1st}$ | 30 | 20 | 20 | −30 | 0.7 |
| $\beta_{2nd}$ | 30 | 20 | 20 | 20 | 0.9 |

**Recovery power stroke** (Equation S38).

| **Function** | $E_{R*}$ $[k_B T]$ | $A_{R*}$ [°] | $B_{R*}$ [°] | $\eta_{R*}$ | $C_{R*}$ |
|---|---|---|---|---|---|
| $\alpha_{RS,RD}$ | −6 | 20 | 20 | −100 | 0.4 |
| $\alpha_{RS,R2}$ | −4 | 32 | 32 | 40 | 0.6 |

### H. Hooking interaction between the myosin head and actin filament

The hooking interaction between the myosin head and actin filament is introduced to prevent sliding of the myosin head along the actin filament. This interaction is assumed to be effective through all states and modeled based on the two variables $r_k$ and $h_k$, which are defined by the binding point $\boldsymbol{x}_0$ in the myosin head and two consecutive points, $\boldsymbol{y}_{0,k}$, $\boldsymbol{y}_{1,k}$, in the actin filament (see Figure 3) as follows.

$$r_k = \|\boldsymbol{x}_0 - \boldsymbol{y}_{0,k}\| \,, h_k = \left(\boldsymbol{x}_0 - \boldsymbol{y}_{0,k}, \frac{\boldsymbol{y}_{1,k} - \boldsymbol{y}_{0,k}}{\|\boldsymbol{y}_{1,k} - \boldsymbol{y}_{0,k}\|}\right). \qquad \text{(S39)}$$

The potential is represented as

$$\varphi_{pas,bind} = \sum_k E_{Hook} \lambda_{A,R_0}(r_k) \mu_{B,H_0}(h_k), \qquad \text{(S40)}$$

with the two functions:

$$\lambda_{A,R_0}(r) = \begin{cases} 0, r < R_0 - A \\ (1 - \cos(\pi(r - R_0 + A)/A))/2\,, R_0 - A \leq r < R_0 + A \\ 0, r \geq R_0 + A \end{cases} \quad \text{(S41)}$$

and

$$\mu_{B,H_0}(h) = \begin{cases} 0, h < H_0 \\ (1 - \cos(\pi(h - H_0)/B))/2\,, H_0 \leq h < H_0 + B, \\ 1, h \geq H_0 + B \end{cases} \quad \text{(S42)}$$

where $E_{Hook}$ = 5 pN · nm, A = 2 nm, $R_0$ = 4 nm, B = 1 nm, and $H_0$= 0 nm.

**Supplementary Movie 1. High-speed AFM movie showing the detachment of myosin from actin.**

The dynamic process at 1 μM caged ATP was filmed at 400 ms $frame^{-1}$ (2.5 frames $s^{-1}$). Image area, 60 × 84 $nm^2$ with 20 × 28 pixels.

**Supplementary Movie 2. High-speed AFM movie showing the two-step lever-arm swing of myosin II.**

The dynamic process at 1 μM caged ATP was filmed at 400 ms $frame^{-1}$ (2.5 frames $s^{-1}$). Orientation changes in myosin II S1 were observed in a two-step manner. Image area, 60 × 84 $nm^2$ with 20 × 28 pixels.

**Supplementary Movie 3. Dynamical process of the numerical high-speed AFM model.**

The transitions of conformational states for 40 s are displayed with the same colors as Figure 2. The individual spirals in the actin filament are displayed blue and red. There are small and large jumps in the transitions from the 2nd post-power stroke state to the pre-power stroke state. The myosin heads land on the same spiral in a short time for small jumps, whereas they land on the other spiral taking a relatively long time for large jumps.

**Supplementary Movie 4. Dynamical process of the numerical sarcomere model with 42.8 nm myosin spacing.**

The transitions of conformational states for 40 s are displayed with the same colors as Figure 6. Two protofilaments in the actin filament are displayed blue and red, respectively. At the right edge of the actin filament (not displayed in the movie), the rotation is prohibited by connecting with the Z-line and a pulling force of 50 pN. The clusters of three or four attached myosin molecules move right to left. The 1st and 2nd post-power stroke transitions also propagate right to left. At the front of each cluster, the jumps to another actin protofilament take place after detachment from the 2nd post-power stroke state.

**Supplementary Movie 5. Dynamical process of the numerical sarcomere model with 37 nm myosin spacing.**

Transitions of conformational states for 40 s are displayed with the same colors as in Figure 7. Two protofilaments in the actin filament are displayed blue and red, respectively. At the right edge of the actin filament, the rotation is prohibited by connecting with the Z-line and a pulling force of 50 pN.
